# Systematic identification of conditionally folded intrinsically disordered regions by AlphaFold2

**DOI:** 10.1101/2022.02.18.481080

**Authors:** T. Reid Alderson, Iva Pritišanac, Đesika Kolarić, Alan M. Moses, Julie D. Forman-Kay

**Author notes:** Equal contribution (T. Reid Alderson, Iva Pritišanac).

## Abstract

The AlphaFold Protein Structure Database contains predicted structures for millions of proteins. For the majority of human proteins that contain intrinsically disordered regions (IDRs), which do not adopt a stable structure, it is generally assumed these regions have low AlphaFold2 confidence scores that reflect low-confidence structural predictions. Here, we show that AlphaFold2 assigns confident structures to nearly 15% of human IDRs. By comparison to experimental NMR data for a subset of IDRs that are known to conditionally fold (i.e., upon binding or under other specific conditions), we find that AlphaFold2 often predicts the structure of the conditionally folded state. Based on databases of IDRs that are known to conditionally fold, we estimate that AlphaFold2 can identify conditionally folding IDRs at a precision as high as 88% at a 10% false positive rate, which is remarkable considering that conditionally folded IDR structures were minimally represented in its training data. We find that human disease mutations are nearly 5-fold enriched in conditionally folded IDRs over IDRs in general, and that up to 80% of IDRs in prokaryotes are predicted to conditionally fold, compared to less than 20% of eukaryotic IDRs. These results indicate that a large majority of IDRs in the proteomes of human and other eukaryotes function in the absence of conditional folding, but the regions that do acquire folds are more sensitive to mutations. We emphasize that the AlphaFold2 predictions do not reveal functionally relevant structural plasticity within IDRs and cannot offer realistic ensemble representations of conditionally folded IDRs.

**Significance Statement:** AlphaFold2 and other machine learning-based methods can accurately predict the structures of most proteins. However, nearly two-thirds of human proteins contain segments that are highly flexible and do not autonomously fold, otherwise known as intrinsically disordered regions (IDRs). In general, IDRs interconvert rapidly between a large number of different conformations, posing a significant problem for protein structure prediction methods that define one or a small number of stable conformations. Here, we found that AlphaFold2 can readily identify structures for a subset of IDRs that fold under certain conditions (conditional folding). We leverage AlphaFold2’s predictions of conditionally folded IDRs to quantify the extent of conditional folding across the tree of life, and to rationalize disease-causing mutations in IDRs.

**Classifications**: Biological Sciences; Biophysics and Computational Biology

## Introduction

The accurate prediction of protein structures from amino-acid sequences has been a long-term goal in biology (Anfinsen 1973; Baker & Sali 2001). Two deep learning-based methods, AlphaFold2 (Jumper et al. 2021a) and RoseTTAFold (Baek et al. 2021), have recently enabled protein structure prediction with high accuracy (AlQuraishi 2021). DeepMind subsequently predicted the structures for 98.5% of proteins in the human proteome (Tunyasuvunakool et al. 2021). Proteome-wide structural predictions from many organisms are publicly available, in collaboration with the European Bioinformatics Institute, via the AlphaFold Protein Structure Database (AFDB) (https://alphafold.ebi.ac.uk/) (Varadi et al. 2021). Access to high-quality structural predictions has paved the way for a multitude of applications in structural biology (Akdel et al. 2022; Burke et al. 2023).

An unexpected effect of the AFDB is that it visually demonstrates the prevalence of intrinsically disordered regions (IDRs). IDRs are predicted to comprise *ca.* 30% of the human proteome; play important cellular roles as interaction hubs in transcription, translation, and signaling (Cumberworth et al. 2013; Dyson & Wright 2005); and are enriched in proteins associated with neurological and other diseases (Uversky et al. 2008). Moreover, it has recently become evident that IDRs contribute to and modulate the formation of many *in vivo* biomolecular condensates via multi-valent interactions that lead to phase separation (Borcherds et al. 2021; Martin & Holehouse 2020). Numerous disease-associated mutations are found in IDRs (Vacic et al. 2012; Wong et al. 2020), including mutations implicated in autism-spectrum disorder (ASD) and cancer (Tsang et al. 2020), and aberrant phase separation involving IDRs has been linked to diseases such as ALS, ASD, and cancer (Alberti & Dormann 2019; Tsang et al. 2020), highlighting the need to understand the structural and biophysical impact of these mutations. At the structural level, IDRs are defined by a lack of stable secondary and tertiary structures and rapid interconversion between different conformations (Van Der Lee et al. 2014; Wright & Dyson 2015). Because of their rapid dynamics, IDRs are not amenable to high-resolution structure determination methods and are frequently removed or not observed in structures determined by X-ray crystallography and cryo-electron microscopy. By contrast, AlphaFold2-generated structural models contain the entire protein sequence, including IDRs (Ruff & Pappu 2021), and one can now visualize predictions for the significant fraction of the proteome that was previously “dark” and unobservable (Bhowmick et al. 2016).

IDRs, however, do not adopt the static structures that are depicted in the AFDB (Ruff & Pappu 2021). Instead, IDRs populate an ensemble of interconverting conformations that depends strongly on the primary structure (Das & Pappu 2013; Mittal et al. 2018; Pietrek et al. 2020), and the properties of these ensembles directly impact on the functions of IDRs (Borcherds et al. 2014; Conicella et al. 2020; Das et al. 2016; González-Foutel et al. 2022; Iešmantavičius et al. 2014; Kim et al. 2021; Maltsev et al. 2012; Milles et al. 2015; Mittag et al. 2008; Sugase et al. 2007; Zosel et al. 2018). However, experimentally determined structural information for IDR conformational ensembles constitutes only a tiny fraction of that available for folded proteins (Lazar et al. 2021; Varadi et al. 2014), and such ensembles are not deposited in the Protein Data Bank (PDB) (Burley & Berman 2021), which stores the high-resolution structures that were mined to train AlphaFold2 (Jumper et al. 2021a) and RoseTTAFold (Baek et al. 2021). Protein structure-prediction programs that make use of available data in the PDB, therefore, will be biased by the relatively few available structures of IDRs, which typically involve those IDRs that fold upon binding to an interaction partner (Smith et al. 2021; Wright & Dyson 2009). The presence of IDR structures in the PDB skews the view of other functional states of IDRs and provides no information for myriad other IDRs that do not fit the “folding-upon-binding” paradigm (Borgia et al. 2018; Fuxreiter 2019; Murthy & Fawzi 2020).

Nuclear magnetic resonance (NMR) spectroscopy is well suited to an ensemble-based structural characterization of IDRs at atomic resolution (Jensen et al. 2014; Konrat 2014; Mittag & Forman-Kay 2007). The intrinsic dynamics of IDRs often lead to long-lived NMR signals that can be exploited to collect high-quality data (Ahmed & Forman-Kay 2022; Malki et al. 2021; Sugase et al. 2007; Theillet et al. 2016). Indeed, a battery of NMR experiments have been applied to probe the conformations of IDRs and residual structure therein (Bertoncini et al. 2005; Dyson & Wright 2021; Eliezer 2007; Kakeshpour et al. 2021; Mantsyzov et al. 2014; Salmon et al. 2010), with dedicated software programs focused on integrating NMR and other biophysical methods to determine ensemble representations of IDPs that best agree with the experimental data (Bottaro et al. 2020; Choy & Forman-Kay 2001; Gomes et al. 2020; Krzeminski et al. 2013; Lincoff et al. 2020; Ozenne et al. 2012; Salmon et al. 2010). However, both the integrative structural biology approach used to determine ensemble representations of IDRs and the NMR-driven determination of residual structure or secondary structure propensity in IDRs are often accessible only to specialists. Such data are not deposited in the PDB, which is used to train and validate deep-learning models, such as AlphaFold2. Finally, because AlphaFold2 was trained on a subset of the PDB that excluded NMR structures (Jumper et al. 2021a), NMR data offer a unique validation metric to assess the accuracy of predicted AlphaFold2 structures in solution, as recently demonstrated (Fowler & Williamson 2022; Robertson et al. 2021; Zweckstetter 2021), and here we provide guidelines for its use, including those not requiring time-consuming sequence-specific resonance assignment.

Here, we show that thousands of IDRs are predicted by AlphaFold2 to be folded with high (70 ≤ x < 90) or very high (≥ 90) predicted local difference distance test (pLDDT) scores (Mariani et al. 2013), which measure the confidence in the predicted structures (Jumper et al. 2021b). We find that, compared to IDRs with low pLDDT scores, the amino-acid sequences of IDRs with high pLDDT scores are enriched in charged and hydrophobic residues, show more positional conservation, and have more alignment matches to sequences in the PDB. However, only 4% of IDR sequences with high pLDDT scores have alignment matches in the PDB, indicating that structural templating is not the reason that AlphaFold2 confidently folds these IDRs. For a subset of IDRs that fold under specific conditions, such as in the presence of binding partners (Wright & Dyson 2009) or following post-translational modification (Bah et al. 2015), and have been extensively characterized by NMR spectroscopy, we find that the AlphaFold2 structures of these IDRs resemble the conformation of the folded state. Moreover, for more than 1,400 IDRs that are known to fold under specific conditions, we observed that the AlphaFold2 confidence scores enable the prediction of conditional folding. This suggests that AlphaFold2 can systematically discover disordered regions that fold upon binding or modification. Therefore, we propose that IDRs with high pLDDT scores may fold in the presence of specific binding partners or following post-translational modifications (PTMs), which we refer to as conditional folding.

Then, we leverage the ability of AlphaFold2 to identify conditionally folded IDRs in two separate applications. First, we compare the per-residue mutational burden within IDRs and find that IDRs with high-confidence AlphaFold2 scores are enriched in mutations relative to IDRs with low-confidence scores. Second, we compare predictions of conditional folding in eukaryotes, bacteria, and archaea and find that prokaryotes show much higher proportions of conditionally folding IDRs, leading us to conclude that a large majority of eukaryotic IDRs function without adoption of structure. Finally, we note that the AFDB does not capture the multiplicity of conformational states accessible to IDRs or their dependence on context (interactions or modification), and hence fails to inform on the plasticity that is crucial to the function of IDRs.

## Results

In this work, we focus on the structural predictions that are available in the AFDB (Varadi et al. 2021), which contains pre-computed AlphaFold2 models that can be easily visualized and downloaded for offline inspection. Moreover, for conditionally folded IDRs we find that the structural predictions from the versions of AlphaFold2 that have been implemented as Jupyter Notebooks on Google Colaboratory (Mirdita et al. 2021; Tunyasuvunakool et al. 2021), are generally of lower quality and do not agree well with AFDB (**Supplementary Figure 1**). As such, we focus henceforth exclusively on structural predictions within the AFDB.

We first analyzed the distribution of per-residue pLDDT scores in the human AFDB (**Figure 1A**). The histogram of pLDDT scores shows a clear bimodal distribution, with the local maxima of each distribution centered around values of 100 and 35 (**Figure 1A**). The majority of residues, accounting for 62.6% of the proteome, have pLDDT scores greater than 70 (**Figure 1A**), which is defined to be the lower threshold for a “confident” score. The remaining 37.4% of residues in the proteome have pLDDT scores below 70 (“low”), while 27.8% of residues have scores below 50 (“very low”). Thus, while a significant percentage of residues have “confident” or “very confident” pLDDT scores, suggesting that the predicted structures of regions are expected to be accurate, there also exists a sizeable fraction of residues that have “low” to “very low” pLDDT scores (**Figure 1A**), indicative of low-accuracy structural regions that should not be interpreted quantitatively.

**Figure 1.**
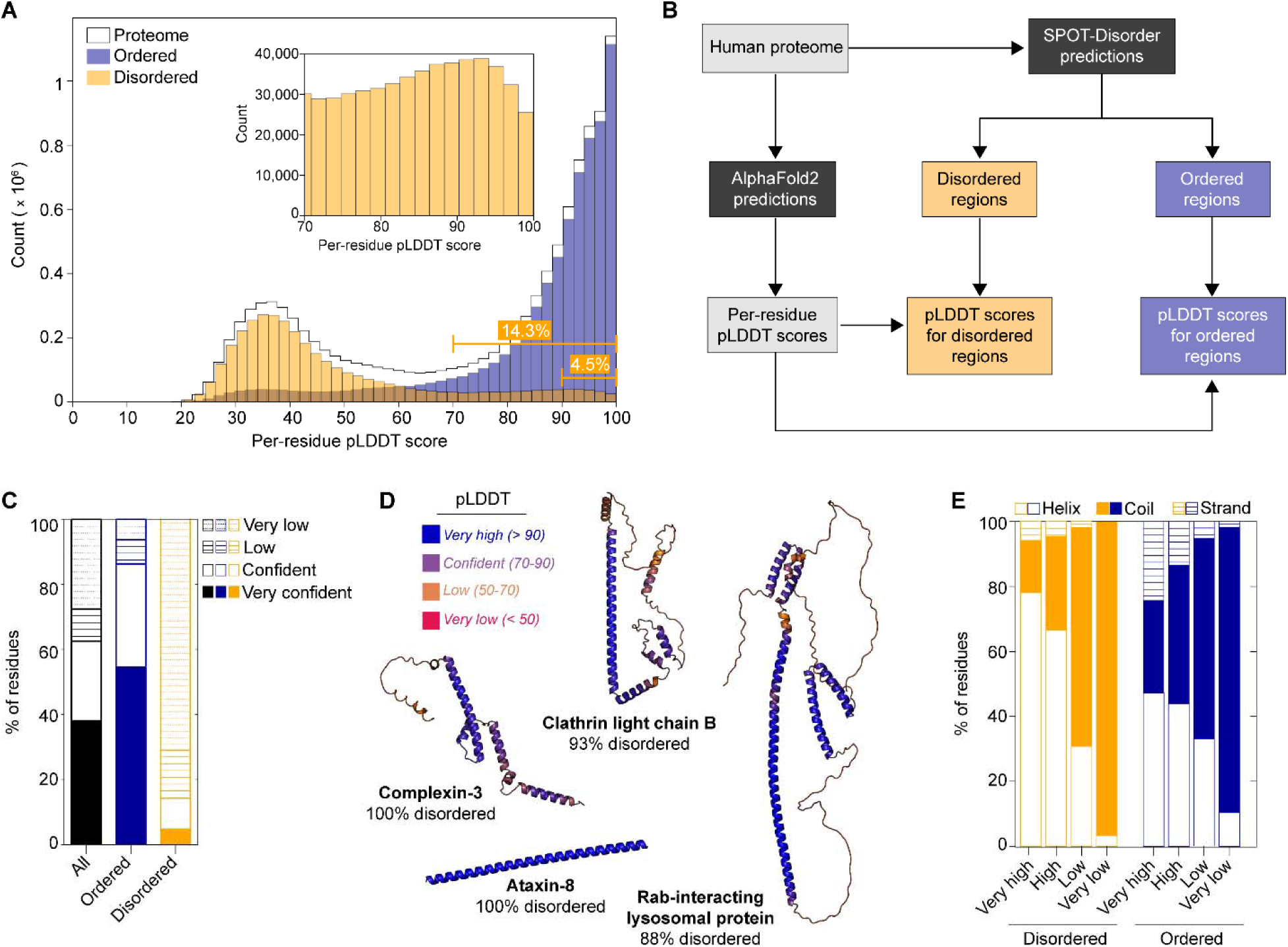
Predicted IDRs in the human proteome that have confident structures in the AFDB. (**A**) Histogram of per-residue pLDDT scores in the human proteome (black) compared with the predicted disordered (orange) or ordered (blue) regions. The inset shows an expansion of the predicted disordered regions between pLDDT scores of 70 to 100. The cumulative percentage of predicted disordered residues with scores greater than or equal to 70 and 90 are indicated in the lower right. (**B**) Flowchart outlining the analysis presented in (A). The human proteome was separated into predicted ordered and disordered regions using a sequence-based predictor of disorder (SPOT-Disorder). Per-residue pLDDT scores were obtained for each protein in the AFDB and split into predicted ordered or disordered regions based on the SPOT-Disorder results. (**C**) Stacked bar graph showing the percentage of residues in the human proteome (black) that have very low (< 50; dotted lines), low (50 ≤ *x* < 70; horizontal lines), confident (≤ 70 *x* < 90; empty), and very confident (≤ 90; filled) pLDDT scores. The corresponding plots are included for SPOT-Disorder-predicted disordered residues (orange) and ordered residues (blue). (**D**) Example structures in the AFDB significantly overlapping SPOT-Disorder-predicted IDRs, with the percentage of predicted disordered residues of the total listed. The AFDB structures have been color-coded by pLDDT scores as indicated. The UniProt IDs are: Q8WVH0 (complexin-3), P09497 (clathrin light chain B), Q96NA2 (Rab-interacting lysosomal protein), and Q156A1 (ataxin-8). (**E**) DSSP-determined secondary structure content of the predicted disordered (orange) and ordered (blue) regions, which were grouped as a function of pLDDT threshold, defined above.

### Predicted human IDRs with high pLDDT scores in the AFDB

To determine how many IDRs in the AFDB have high pLDDT scores that reflect high confidence structural predictions, we extracted the predicted IDRs from the human AFDB (**Figure 1B**). We used the state-of-the-art sequence-based predictor of intrinsic disorder, SPOT-Disorder (Hanson et al. 2017), to calculate the predicted disorder propensities for each protein in the human proteome (**Figure 1B**). A total of *ca.* 3.5 million residues are predicted to be disordered, totalling 32.8% of the human proteome, consistent with expectations based on previous reports (Necci et al. 2021) (**Supplementary Table 1**). We then investigated the proteins that comprise the lower end of the distribution of pLDDT scores within the AFDB. An inverse correlation between pLDDT scores and predicted disorder was previously noted (Tunyasuvunakool et al. 2021), with pLDDT scores reported to perform well as a new predictor of intrinsic disorder (Necci et al. 2021; Tunyasuvunakool et al. 2021). Thus, one might assume that the 32.8% of predicted residues in IDRs would be embedded in the 37.4% of residues with pLDDT scores below 70, because IDRs should have “low” or “very low” confidence structural predictions.

However, when we isolated the pLDDT scores from residues that localize to SPOT-Disorder predicted IDRs (**Figure 1B**), we found that IDRs also have a bimodal distribution of pLDDT scores (**Figure 1A**). Of the *ca.* 3.5 million predicted disordered residues, 14.3% (*i.e.*, ca. 500,000 residues in total) have “confident” pLDDT scores greater than or equal to 70 (**Figure 1A**, **Figure 1C**, **Supplementary Table 2**). When the pLDDT threshold is increased to greater than or equal to 90 (“very confident”), there are more than 160,000 residues that remain, accounting for 4.5% of the total number of disordered residues (**Figure 1A**, **Figure 1C**). This analysis indicates that there is a significant fraction of SPOT-Disorder-predicted IDRs in the human AFDB that have high-confidence structural predictions (**Figure 1A**, **Supplementary Figure 2**), and therefore the assumption that all IDRs have low pLDDT scores is incorrect.

Because the prevalence of confidently predicted structures within IDRs was unexpected, we sought to ensure that the “confident” and “very confident” scores associated with SPOT-Disorder-predicted IDRs are not the result of poor or biased disorder predictions. We extracted the pLDDT scores for IDRs in the DisProt database of experimentally validated IDRs (Quaglia et al. 2021). There is a total of 932 human IDRs in DisProt, yielding over 300,000 residues that can be used as a direct comparison to the SPOT-Disorder predictions. We find that the distribution of pLDDT scores for IDRs in DisProt shows an even higher proportion of residues with “confident” scores greater than or equal to 70 than the SPOT-Disorder-predicted IDRs: nearly 30% of the experimentally validated DisProt IDRs have confident pLDDT scores (**Supplementary Figure 2**). Thus, the IDRs that were predicted by SPOT-Disorder do not contain an artificially inflated fraction of residues with high pLDDT scores. In addition, we checked if the high pLDDT scores might originate from a few disordered residues that are immediately adjacent to structured domains. To this end, we filtered the predicted IDRs with confident pLDDT scores (greater than or equal to 70) for those with consecutive regions of disorder. We found that over 50% of IDRs with high pLDDT scores come from stretches of 24 or more consecutive disordered residues, with nearly 10% arising from IDRs that have 100 or more consecutive disordered residues (**Supplementary Figure 2**). Finally, we filtered the list of IDRs to extract those that have 10 and 30 or more consecutive residues with “confident” (“very confident”) pLDDT scores. We identified 14,996 (4,883) and 3,730 (1,157) IDRs that respectively match these criteria (**Supplementary Table 3**).

From the list of proteins that contain predicted IDRs equal to or longer than 30 residues with high pLDDT scores, we selected a handful of examples for structural analysis in the AFDB (**Figure 1D**). In particular, we identified proteins that are predicted to be predominantly disordered by SPOT-Disorder yet are assigned very high pLDDT scores in the AFDB. For example, the protein ataxin-8 (UniProt ID: Q156A1) contains an initial Met residue followed by 79 Gln residues and is predicted to be fully disordered (**Figure 1D**). However, the AlphaFold2 model indicates that ataxin-8 forms a single α-helical structure with pLDDT scores greater than 90 for every residue in the helix (**Figure 1D**). Similarly, the proteins complexin-3 (UniProt ID: Q8WVH0), clathrin light chain B (UniProt ID: P09497), and Rab-interacting lysosomal protein (RILP, UniProt ID: Q96NA2) all adopt highly α-helical structures with very high pLDDT scores and various degrees of tertiary interactions (**Figure 1D**), despite being predicted by SPOT-Disorder to be almost entirely disordered (88% to 100% predicted disorder).

Given that the above examples were α-helical structures, we computed the secondary structure content for every model in the AFDB to assess the structural properties of IDRs with high-confidence pLDDT scores. This analysis revealed primarily helical conformations in the “high” and “very high” confidence IDR structures (**Figure 1E**). When compared to ordered regions, the predicted IDRs are significantly enriched in helical conformations at the expense of coils and strands (**Figure 1E**, **Supplementary Table 4**). In addition, we note that the predicted IDRs with low confidence scores still exhibit significant secondary structure content: over 32% of residues with pLDDT scores between 50 and 70 are assigned to regions of secondary structure as compared to 38% in ordered regions (**Figure 1E**, **Supplementary Table 4**). In the IDRs with pLDDT scores < 50, the percentage of residues in regions of secondary structure dramatically diminishes to only 3.4% (**Figure 1E**, **Supplementary Table 4**).

Overall, the analysis of secondary structure content in predicted IDRs with high pLDDT scores shows an enrichment in helical conformations. Moreover, the selected examples in **Figure 1D** all have at least one long, extended α-helix that is not stabilized by tertiary contacts. These so-called single α-helix (SAH) domains are well known in the literature and are estimated to exist in 0.2-1.5% of human proteins (Barnes et al. 2019; Swanson & Sivaramakrishnan 2014), with the formation of SAHs dependent on stabilizing *i* to *i*+4 salt bridges between charged side chains (Marqusee & Baldwin 1987). SAH domains are enriched in E, K, and R residues, which form in various repeats and sum to nearly 80% of the residues in SAH sequences, while the remaining 20% of the sequences derive from A, D, L, M, and Q residues, (Simm & Kollmar 2018). The long α-helix in ataxin-8, which is composed almost entirely of Q, represents another form of an experimentally observed SAH. NMR analyses recently showed that regions containing homo-repeats of Q (polyQ) also form SAHs via side-chain *i* to main-chain *i*-4 hydrogen bonds, although the helical structure decays toward the C-terminal region of the polyQ tract and is dependent on non-Gln hydrogen-bond acceptors at the N-terminus of the helix (Escobedo et al. 2019). Thus, although the sequences of the protein regions in **Figure 1D** would not be classified as *bona fide* SAH domains, SAH-like structures in the AFDB for sequences predicted to be IDRs may be plausible and physically reasonable. It is likely that these SAH-like structural regions for predicted IDRs in the AFDB represent structural states that are stabilized upon interactions or modification, including a combination of canonical SAHs and long α-helices that form stabilizing inter-molecular contacts (*e.g.*, coiled coils).

### Comparing NMR data and AlphaFold2 structures for experimentally characterized IDRs

Given that our structural analyses above relied on sequence-based predictions of intrinsic disorder, we asked whether the structures of IDRs with high pLDDT scores show correspondence with experimentally determined structural propensities of IDRs. To this end, we focus on three IDRs/IDPs that have been characterized in detail by NMR spectroscopy (Alderson et al. 2018; Bah et al. 2015; Bodner et al. 2009; Dawson et al. 2020; Demarest et al. 2002; Ebert et al. 2008; Eliezer et al. 2001; Mantsyzov et al. 2015; Marsh et al. 2006; Salmon et al. 2010; Theillet et al. 2016). Two of the model proteins, α-synuclein (UniProt ID: P37840) and 4E-BP2 (UniProt ID: Q13542), are full-length IDPs, whereas the third protein, ACTR or NCoA3 (UniProt ID: Q9Y6Q9), is a small IDR that is part of a larger protein with folded domains and other longer IDRs. The AlphaFold2-predicted structures of the three proteins (**Figure 2A**) vary from all helical (α-synuclein, ACTR) to a mixture of strand and helix (4E-BP2). For each structure, the pLDDT scores in the regions of secondary structure range from “high” to “very high” (**Figure 2B**), suggestive of atomic-level accuracy and an overall high level of confidence in the structural models (Jumper et al. 2021a; Varadi et al. 2021). Next, we checked the predicted disorder propensity using four different sequence-based predictors of intrinsic disorder (Emenecker et al. 2021; Hanson et al. 2017; Jones & Cozzetto 2015; Mészáros et al. 2018), and we found that either two (4E-BP2, ACTR) or three (α-synuclein) of the four programs predicted that these proteins would be predominantly ordered (**Figure 2C**). Thus, without additional experimental evidence, an AFDB user who relies on the overlap between sequence-based disorder prediction software and the (confident) AFDB structure would likely assume that the IDR/IDP under investigation folds into the high-confidence predicted structure.

**Figure 2.**
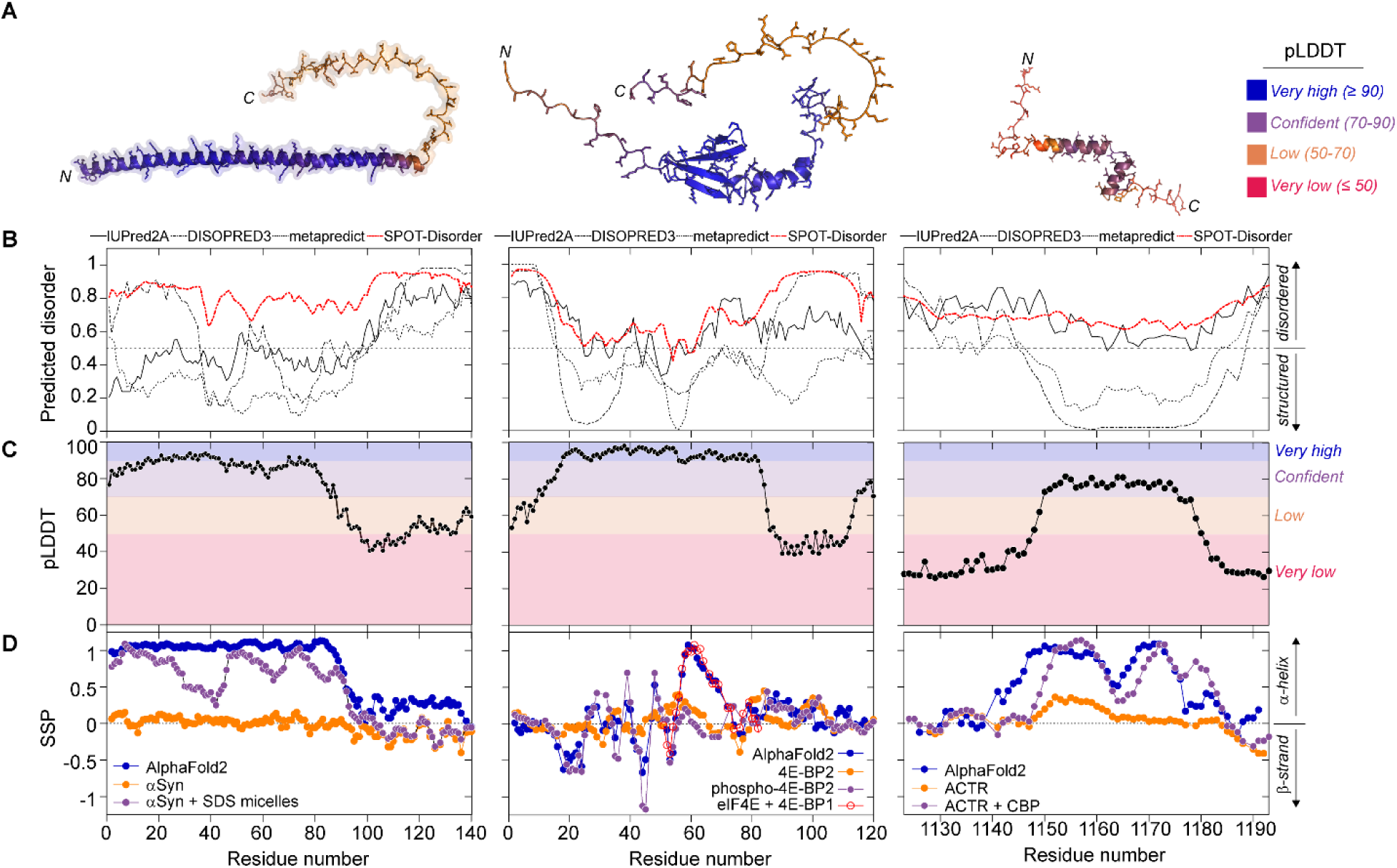
Examples of three IDRs with high pLDDT scores that conditionally fold and have been characterized by NMR spectroscopy. **(A)** AlphaFold2-predicted structures of three IDPs/IDRs that have been extensively characterized by NMR spectroscopy. The structures are color coded by pLDDT scores. From left to right: α-synuclein, 4E-BP2, and ACTR. The N- and C-termini of each protein are indicated. **(B)** Sequence-based prediction of disorder for the three IDPs/IDRs. Four different programs were used: IUPred2A, DISOPRED3, metapredict, and SPOT-Disorder. Only SPOT-Disorder correctly predicts the disordered nature of all three IDPs/IDRs. **(C)** Per-residue pLDDT confidence scores derived from the AlphaFold2 structures. The thresholds are defined according to the AFDB: very high (90-100), confident (70-90), low (50-70), and very low (< 50). **(D)** NMR ^13^Cα and ^13^Cβ chemical shift-derived secondary structure propensity (SSP) (Marsh et al. 2006). For the AlphaFold2 structures (blue) and the 4E-BP2 peptide bound to eIF4E (PDB ID: 3am7), NMR chemical shifts were back-calculated from the structure using SPARTA+ software (Shen & Bax 2010). The unbound/unmodified IDPs/IDRs (orange) show very little preferential secondary structure (α-synuclein) or modest populations of helix (4E-BP2, ACTR). By contrast, the binding to SDS micelles (α-synuclein), phosphorylation (4E-BP2), binding to eIF4E (4E-BP2), or the binding to CBP (ACTR) induces the formation of stable secondary structure (purple) that is in better agreement with the AlphaFold2 structures.

However, we find that there is disagreement between experimental NMR data from these IDRs/IDPs and the AlphaFold2 models and sequence-based prediction of disorder (**Figure 2D**). It is well known that ^13^Cα and ^13^Cβ chemical shifts are sensitive reporters on the secondary structure of a protein (Cornilescu et al. 1999; Spera & Bax 1991; Wishart et al. 1991). For each residue in the protein, the expected chemical shifts for a fully disordered state can be subtracted from the measured chemical shifts. These so-called secondary chemical shifts (corrected for neighboring residues) provide residue-level information regarding the secondary structure of a protein, including the fluctuating, fractional secondary structure of disordered regions, which can be quantified using software programs such as δ2D, SSP, and CheSPI (Camilloni et al. 2012; Marsh et al. 2006; Nielsen & Mulder 2021). If the AFDB structures were correct, the expected secondary chemical shifts would reveal long stretches of secondary structure with “fractional” structure values near 1 or -1 for a fully stabilized α-helix or a β-strand, respectively (**Figure 2D**). By contrast, the experimental NMR data for each of three proteins in **Figure 2** show that there is no stable secondary structure and only a fractional preference to populate secondary structure (**Figure 2D**). Thus, a user without knowledge of the disordered nature of these proteins from experiment could erroneously trust the confident AlphaFold2 models (**Figure 2A**, **Figure 2C**) and use the lack of predicted disorder (**Figure 2B**) as a cross-validation method to justify the structures.

Interestingly, however, there are correlations between the AFDB structures of these IDRs/IDPs and their experimentally defined conformations under specific conditions. For example, the N-terminal *ca.* 100 residues of α-synuclein fold into a long α-helix in the presence of lipid vesicles (Bodner et al. 2009), and the AFDB structure reflects this lipid-bound conformation (**Figure 2A**). There is no high-resolution structure of the lipid-bound state, so no side-by-side structural comparison can be made; assigned NMR chemical shifts are only available for α-synuclein bound to SDS micelles (Chandra et al. 2003) (**Figure 2D**, purple). The solution structure of 4E-BP2 revealed that it folds into a four β-strand structure upon multi-site phosphorylation (PDB ID: 2mx4) (Bah et al. 2015), and the AFDB structure has correctly identified the β-strands and the intermolecular contacts: the heavy-atom RMSD is 0.35 Å upon alignment of the β-strands from T19-D55 in the experimental structure to the AFDB model. Confusingly, however, an additional helix in the AFDB model is present in residues R56-R62, followed by a short turn and then a 3_10_-helix between residues P66-Q69. These additional helices resemble those seen in crystal structures of fragments of non-phosphorylated 4E-BP2 and 4E-BP1 bound to eukaryotic translation initiation factor 4E (eIF4E) (PDB IDs: 3am7, 5bxv). For ACTR, a helix-turn-helix motif is present in the AFDB structure, whereas a three-helix structure is formed upon binding to CBP (PDB: 1kbh). ACTR and 4E-BP2 provide particularly useful test cases because the experimental structures (PDB ID: 1kbh, 2mx4) were determined by NMR spectroscopy, and AlphaFold2 was not trained on NMR structures (Jumper et al. 2021a).

Finally, we note that fractional secondary structure in the unbound form of an IDR does not necessarily correlate with the secondary structure in the AFDB model of the IDR. For example, in the unbound form of ACTR, there is an α-helix populated to approximately 40% between residues 1150-1163 (**Figure 2D**, orange), which closely matches the position of the first α-helix in the AFDB (**Figure 2D**, blue) and experimental structures (**Figure 2D**, purple). However, the second and third α-helices in the experimental structure are not appreciably formed in the unbound state or the AFDB model (**Figure 2D**, orange). The lack of clear correlation between fractional secondary structure in an isolated IDR and stabilized structure in complexes has been noted previously (Marsh et al. 2010).

These comparisons show that the high-confidence AFDB structures of IDRs do not reflect the conformational ensemble sampled by the unbound or unmodified form of the IDR. Instead, the AlphaFold2 structures appear to resemble conditionally folded states. Moreover, in the case of 4E-BP2, the AlphaFold2 model has combined structural features from two different conditionally folded forms of the protein that do not coexist: one structure forms upon multi-site phosphorylation (β-strand-rich) and the other upon binding to eIF4E (helical). In this case, the AlphaFold2 structure of 4E-BP2 obscures the molecular mechanism of the protein function (see section below).

### AlphaFold2 structures of experimentally characterized IDRs resemble the conditionally folded state but do not capture structural plasticity

Our analysis of IDRs/IDPs with extensive NMR data showed that AFDB models with high pLDDT scores might reflect a conformation of the IDR/IDP that only forms under specific conditions. We thus examined the AlphaFold2 structures of an additional four IDRs/IDPs that are known to fold upon binding to interacting partners and have high-resolution structures of the complex in the PDB. The structures for two of these complexes were determined by X-ray crystallography (p27: 1jsu; SNAP25: 1kil) and two were determined by NMR spectroscopy (HIF-1α: 1l8c; CITED2: 1p4q). A comparison of the experimental structures (**Figure 3A-D**) with those in the AFDB (**Figure 3E-H**) shows an overall high structural similarity (**Figure 3I-L**). For the two examples with very confident AFDB structures (p27, SNAP25), the heavy atom root-mean-squared-deviations (RMSDs) when comparing the experimental and the AFDB structures are 0.5 and 2.1 Å (**Figure 3E,F,I,J**). Even for some structures that have a mixture of very confident and low pLDDT scores (CITED2, **Figure 3F, 3G**), or only low pLDDT scores (HIF-1α, **Figure 3H**), the overall architecture of the AFDB structure resembles that of the experimental structure, with RMSD values of 1.6 (**Figure 3K**) and 5.0 Å (**Figure 3L**), respectively. Taken together, these analyses suggest that the AFDB structures formed by IDRs with high pLDDT scores are likely capturing some structural features that form in the presence of specific interactions. In the case of an IDR with very low pLDDT scores (HIF-1α), the regions of secondary structure appear to correlate with the bound-state conformation.

**Figure 3.**
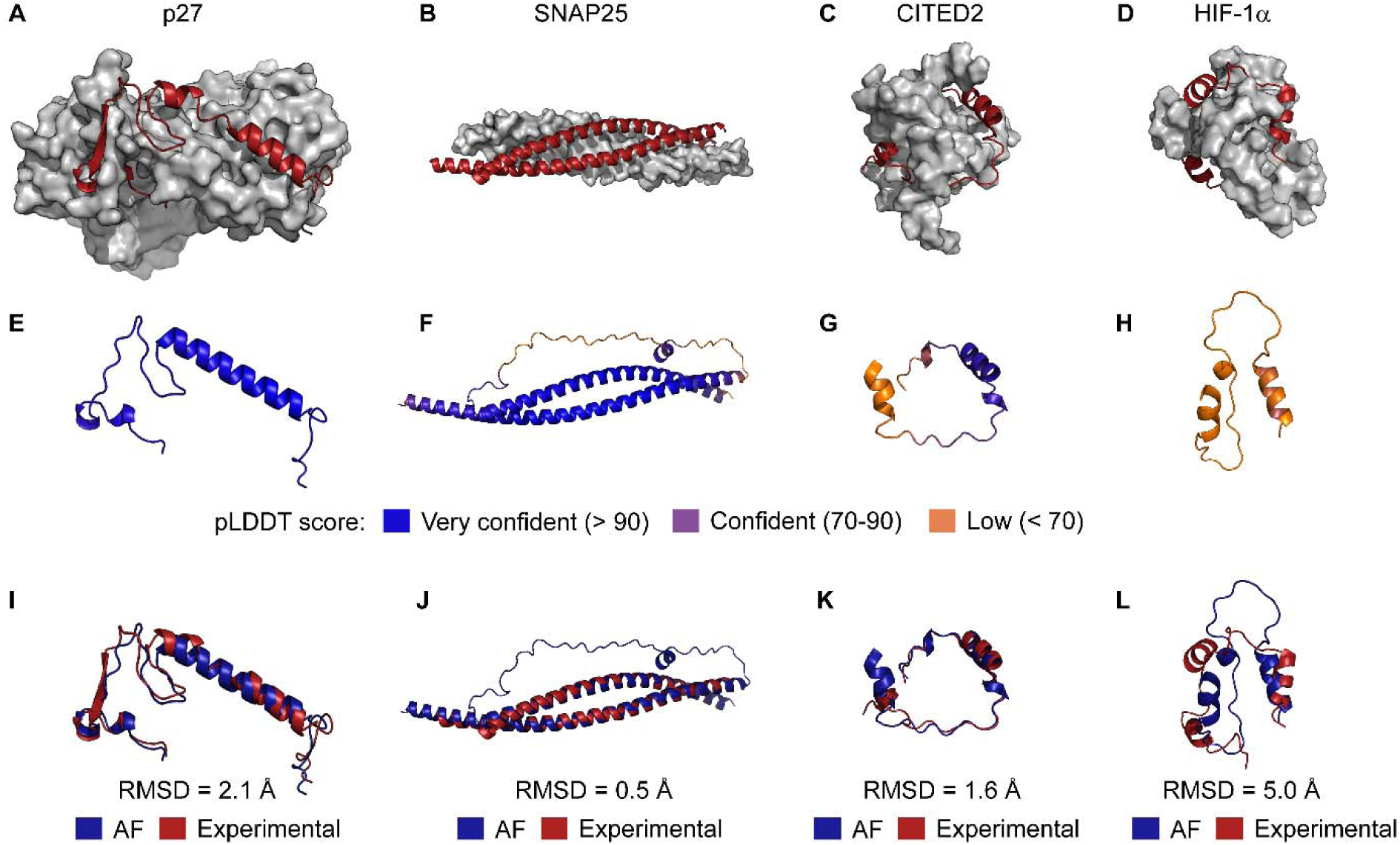
Structures of IDRs in the AFDB correlate with experimentally determined structures of the IDRs bound to interaction partners. **(A, B, C, D)** experimental structures for the listed IDRs/IDPs (red) bound to an interacting folded domain (grey surface representation). The PDB ID codes are 1jsu, 1kil, 1p4q, and 1l8c respectively. **(E, F, G, H)** the predicted structures in the AFDB for the listed IDRs/IDPs in panels A-D. These structures have been color-coded by per-residue pLDDT scores, with blue, purple, and orange respectively corresponding to very confident (≥ 90), confident (70-90), and low (< 70) scores. **(I, J, K, L)** comparison of the experimental structures from panels A-D with the predicted structures in the AFDB from panels E-H. Experimental structures are colored red and AFDB structures blue. The heavy-atom RMSD upon alignment of secondary structure elements is indicated.

If AlphaFold2 structures of IDRs reflect the conditionally folded state, we were interested to determine if these structures can inform on the molecular mechanisms of IDRs. The interconversion between different structural forms is essential for IDR function, and it is well known that IDRs can bind to multiple interaction partners via different interfaces or motifs, often times forming unique structural elements in the process (Wright & Dyson 2009). We selected three IDRs with multiple experimentally determined structures in which the IDR has folded into a different conformation. Some of the experimental structures of these IDPs or IDRs from the cystic fibrosis transmembrane conductance regulator (CFTR; UniProt ID: P13569), SNAP-25, and 4E-BP2 show good agreement with the AlphaFold2 models (**Figure 4A-C**). For example, the regulatory (R) region of CFTR is a long IDR that is heavily phosphorylated with several regions that adopt residual helical propensity (Baker et al. 2007; Bozoky et al. 2013b). The weak, multivalent inter- and intramolecular interactions between the R region and different binding partners regulate the activity of CFTR in a phosphorylation-dependent manner (Bozoky et al. 2013b). The AlphaFold2 model of the portion of the CFTR R region immediately following the first nucleotide-binding domain (NBD1), called the regulatory extension (RE), shows close agreement with its conformation in a crystal structure of NBD1 and the RE (**Figure 4D**). However, another structure of NBD1 shows the RE interacting with a different interface on NBD1, with the orientation of RE with respect to NBD1 dramatically altered, despite almost no changes to the structure of NBD1 itself (Bozoky et al. 2013a). The AFDB structure of CFTR contains only one of these conformational states for the RE.

**Figure 4.**
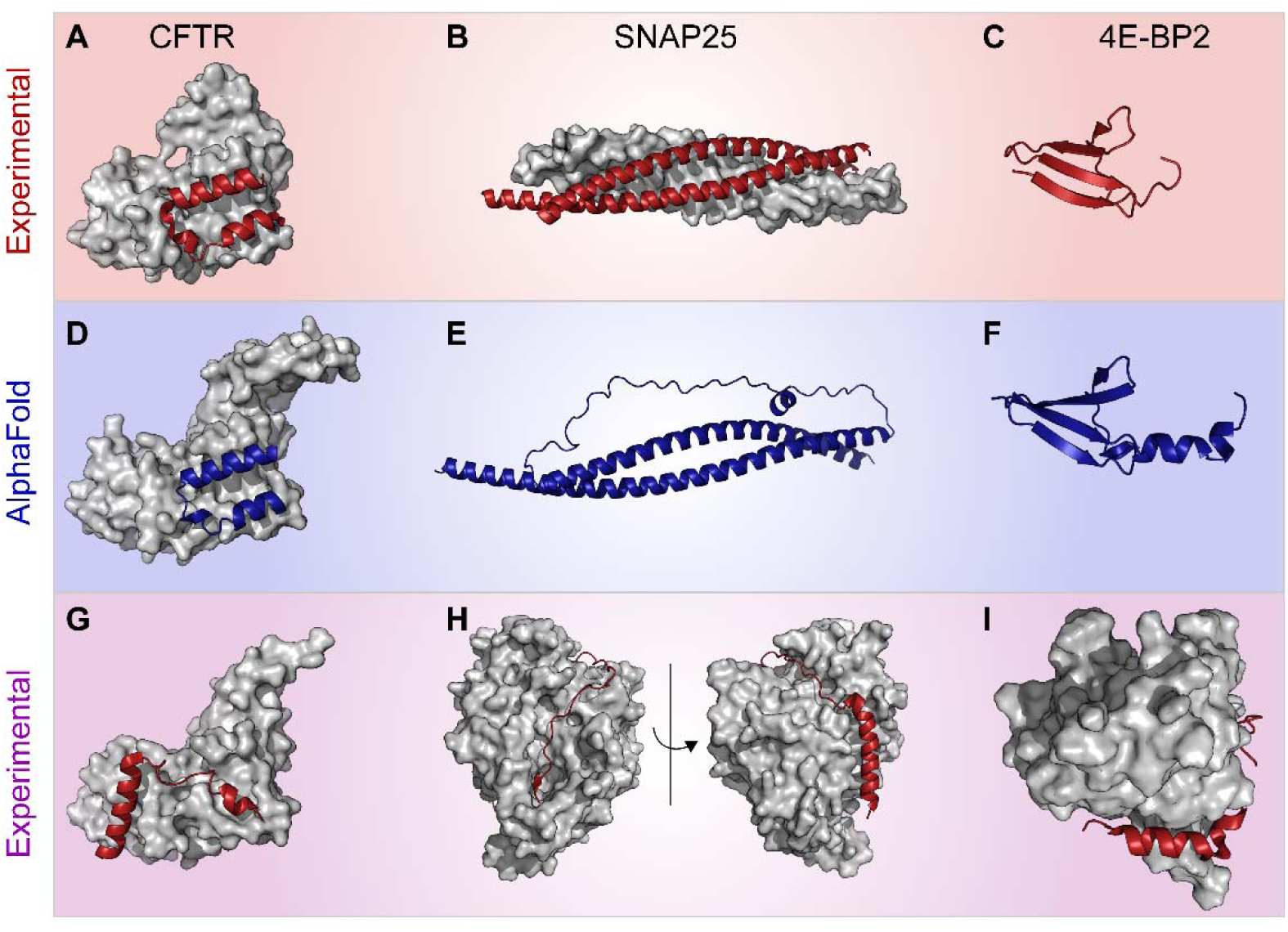
The AFDB does not capture the inherent structural plasticity of IDRs/IDPs. Shown here (top) are three examples of IDRs/IDPs that have experimentally determined structures when bound to interacting partners. (**A**) The disordered regulatory extension (RE) of human CFTR (residues L636-S670) bound intramolecularly to NBD1 (residues S388-S635; PDB ID: 1r0x). Note that the N-terminal region of NBD1 contains a cloning artifact and so residues S388-I393 differ from the AlphaFold2 model. (**B**) The N- and C-terminal SNARE motifs of human SNAP25 (residues S10-L81 and G139-W204, respectively) bound to rat VAMP2 (residues S28-N93), rat syntaxin-1A (residues L192-D250), and rat complexin-1 (residues K32-I72; PDB ID: 1kil). (**C**) Phosphorylated 4E-BP2 (residues P18-R62; PDB ID: 2mx4. (**D**-**F**) The AlphaFold2-predicted structures (blue) of the CFTR RE (residues P638-S670), SNAP25 (M1-G206), and 4E-BP2 (A16-P72) show excellent correspondence with the experimental structures in A-C. (**G**-**I**) However, the CFTR RE, SNAP25, and 4E-BP2 have also been captured in a different conformation as those in panels A-C (red). (**G**) CFTR RE (P638-L671) intramolecularly bound in a different orientation to NBD1 (S388-Q637; PDB ID: 1xmi). (**H**) SNAP25 C-terminal SNARE motif (residues M146-G204) bound to BoNT/A) protease (residues P2-R425; PDB ID: 1xtg). (**I**) A peptide from 4E-BP2 (residues T50-P84) bound to eIF4E (residues H33-K206; PDB ID: 5bxv).

Another example is provided by SNAP-25, which is an IDP that folds into a helical bundle in SNARE complexes (**Figure 4B**) that have important functions in membrane fusion during synaptic vesicle exocytosis. The AlphaFold2 model of SNAP-25 correctly identifies the N- and C-terminal soluble N-ethylmaleimide-sensitive factor attachment protein receptor (SNARE) motifs that form a four-helix bundle in the ternary SNARE complex involving SNAP-25, synaptobrevin, and syntaxin (**Figure 4E**). However, SNAP-25 is also a substrate for Clostridal neurotoxins (CNTs), which are zinc-dependent endopeptidases that cause the diseases tetanus and botulism by specifically cleaving SNARE proteins and impairing neuronal exocytosis (Schiavo et al. 1992). The crystal structure of the botulinum neurotoxin serotype A (BoNT/A) protease bound to SNAP-25 reveals an extensive interface involving the C-terminal SNARE motif of SNAP-25 (Breidenbach & Brunger 2004) (**Figure 4I**). An α-helix is formed in BoNT/A-bound SNAP25 by residues D147-M167, while the remaining residues G168-G204 of SNAP-25 are bound to BoNT/A in a coil conformation, with a small β-strand (K201-L203) formed as well (Breidenbach & Brunger 2004) (**Figure 4I**). By contrast, in the SNARE complex bound to complexin-1, SNAP25 forms a long α-helix that encompasses residues S140-M202 (Chen et al. 2002) (**Figure 4B**). These three structures of SNAP-25 provide an illustrative example: given the very high confidence in the AlphaFold2 structure of SNAP-25, one could assume that an ordered-to-disordered transition is required for SNAP-25 to bind to BoNT/A in the experimentally observed conformation, and that SNAP-25 assembles into SNARE complexes as a rigid body with minimal structural changes. Both of these assumptions are in stark contrast to the known disordered-to-ordered transition that occurs both upon binding to BoNT/A and formation of the SNARE complex. Thus, the molecular mechanism of SNAP-25 function, and its proteolytic cleavage in disease, are obscured by the high-confidence AlphaFold2 model.

Finally, we examined the case of 4E-BP2, an IDP that is a regulatory binding protein for the eIF4E, with experimental structures of segments of the protein in the 5-site phosphorylated state and in the non-phosphorylated state bound to eIF4E (**Figure 4C, 4F, 4J**), as discussed in the section above. The AlphaFold2 structure of residues A16-P72 of 4E-BP2 contains a β-sheet followed by α- and 3_10_-helices (**Figure 4F**), whereas the experimental structure in phosphorylated 4E-BP2 contains the β-sheet for residues T19-D55 followed by a coil region (Bah et al. 2015) (**Figure 4C**). The AlphaFold2 model correctly places the β-strands in phosphorylated 4E-BP2 and accurately identifies the orientations of each strand relative to one another (**Figure 4F**). However, the helical secondary structure elements are only observed when non-phosphorylated 4E-BP2 binds to eIF4E (PDB ID: 3am7) (**Figure 4J**). Upon binding to eIF4E, a truncated peptide from 4E-BP2 was shown to fold into an α-helix between residues D55-D61 (**Figure 4J**). The related protein 4E-BP1, for which more complete structural information is available when bound to eIF4E (Sekiyama et al. 2015) (PDB ID: 5bxv), forms an α-helix between residues D55-R62 followed by a short turn and a 3_10_-helix between residues P66-Q69. The phosphorylation-induced folded state inhibits the binding of eIF4E that is otherwise extremely tight for the unmodified protein (Bah et al. 2015). The β-strand-rich structure of the AlphaFold2 model of 4E-BP2 is incompatible with the binding to eIF4E, which requires the disordered non-phosphorylated state known to fractionally sample helical structure in the segment that forms a stable α-helix in complex with eIF4E (Lukhele et al. 2013). Thus, the 4E-BP2 AFDB structure reflects a strange mixture of the ordered landscape of the protein, combining that found in the presence of PTMs with that stabilized in the absence of PTMs but in the presence of a protein binding partner, and confounding understanding of the mechanism of phospho-regulation of translation initiation (Bah et al. 2015).

### Rapid experimental assessment of the accuracy of AlphaFold2 predictions

Given that the AFDB contains only a single structure of a given IDR, our analyses above emphasize that these AlphaFold2 models are but one possible conformation of the IDR, especially in the context of an IDR binding to an interaction partner. Moreover, because predicted IDRs/IDPs with high or very high pLDDT scores are more widespread than initially thought (**Figure 1A**), it is critical to be able to determine if the AFDB structures are correct. The comparisons outlined above relied on known structural knowledge available, NMR data with assigned chemical shifts enable an atomic-level comparison of the AlphaFold2-predicted structure and the protein conformations that are present in solution (Fowler & Williamson 2022; Gelenter & Bax 2023; Robertson et al. 2021; Zweckstetter 2021). However, the assignment of NMR chemical shifts requires considerable time and effort.

In the absence of chemical shift assignments, an integrative biophysical approach that requires only limited amounts of purified, unmodified protein can rapidly assess the accuracy of AlphaFold2 predictions for IDRs (**Figure 5A, 5B, 5C**). We demonstrate this approach using the IDP α-synuclein as a model system, for which many biophysical measurements are available. Whereas the AlphaFold2 model of α-synuclein is highly helical, the circular dichroism (CD) spectrum of the protein in solution shows that it is largely devoid of global secondary structure content (**Figure 5A**). The molecular dimensions of α-synuclein, derived from pulsed-field gradient NMR (PFG-NMR) translational diffusion and small-angle X-ray scattering (SAXS) data, namely the gyration and hydration radii (*R*_g_ and *R*_h_) and the maximum interatomic distance (*D*_max_), are also significantly smaller than the predicted model (**Figure 5B, 5C**), suggesting a more compact ensemble of states is present in solution.

**Figure 5.**
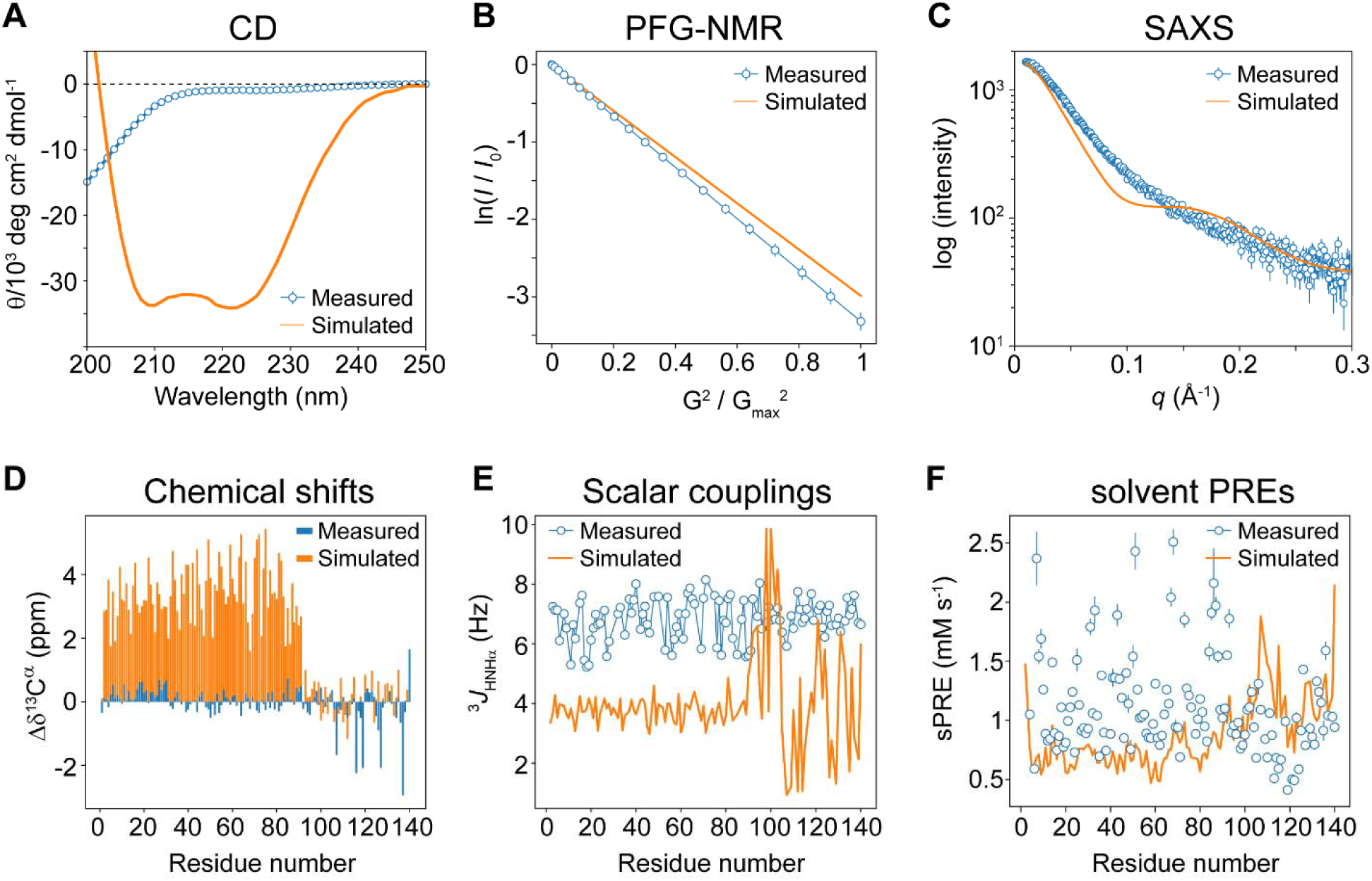
Integrative biophysical approach to evaluate high-confidence structures of IDRs in the AFDB. Biophysical experiments can rapidly determine if an AFDB structure accurately captures the global properties of the protein in solution. Experimental (**A**) CD spectrum (Fusco et al. 2016), (**B**) PFG-NMR translational diffusion (Ramanujam et al. 2020a), and (**C**) small-angle X-ray scattering (SAXS) data (Ahmed et al. 2021) for α-synuclein are shown in blue. Simulated data based on the AFDB structure of α-synuclein are shown in orange. These experiments require a limited amount of unmodified, natural abundance protein samples. If NMR assignments are available, local structural details can be compared, although such experiments require considerably more time as well as isotope-enriched samples. Experimental data for α-synuclein are shown in blue. (**D**) secondary ^13^Cα chemical shifts (Bermel et al. 2006), (**E**) ^3^*J*_HNH_ coupling constants (Mantsyzov et al. 2014), and (**F**) ^1^Hα solvent paramagnetic relaxation enhancements (PREs) (Hartlmüller et al. 2019). Even if NMR assignments are not available, the data shown in **D**-**F** can be plotted as histograms to compare experimental results with the simulations. The secondary ^13^Cα shifts can be recomputed as ^13^Cα – ^13^Cβ shifts, because the secondary shifts require knowledge of the random coil shifts, which would not be known without assignments. See the Supplementary Appendix for more details.

If NMR assignments are available, then atomic-level structural details in solution can be probed, such as dihedral angles and local solvent accessibility. Indeed, the experimental secondary ^13^Cα chemical shifts (Δδ^13^Cα) and three-bond ^1^H_N_ – ^1^H scalar coupling constants (^3^*J*), which are sensitive to either the φ (^3^*J*_HNH_) or both the φ and ψ dihedral angles (Δδ^13^Cα), deviate significantly from the predicted model (**Figure 5D, 5E**). The experimental ^3^*J*_HNH_ values are largely near 7 Hz, which is consistent with a random-coil conformation. Finally, the solvent paramagnetic relaxation enhancements (sPREs) of α-synuclein reveal that the protein is significantly less protected from solvent than the AlphaFold2 model (**Figure 5F**). Importantly, the data in **Figure 5D-F** are shown with resonance assignments (*i.e.*, per-residue). If assignments are not available, then histograms of the raw values from unassigned peaks can still be used to perform a comparison with the simulated values (**Supplementary Figure 3**).

### Rigid-body docking with confidently AlphaFold2-predicted IDP/IDR structures

The above examples collectively demonstrate how high-confidence AlphaFold2 structures of IDRs/IDPs, which can offer insight into various structures that are accessible to the IDR/IDP, may also obscure the molecular mechanisms of these disordered regions. Next, we investigated this problem from the other side: if high-confidence structures of IDRs/IDPs are capturing the bound/modified states of IDRs/IDPs, then can such structures be used with protein-protein docking software to obtain structural models of IDRs/IDPs could be used with the goal of identifying the interfaces of IDR/IDP-globular domain complexes. As we have shown above (**Figure 2**, **Figure 3**, **Figure 4**), for predicted IDRs/IDPs with high pLDDT scores, AlphaFold2 appears to capture the conditionally folded states of such IDPs/IDRs. Therefore, if the AFDB structure of the IDR/IDP is already in a bound-state conformation, it may be beneficial to use the AlphaFold2 model of the IDR/IDP as a starting structure for a protein-protein docking analysis with a known or putative globular domain interactor. Molecular docking with IDRs/IDPs could be valuable for understanding molecular mechanisms, interaction sites, and binding interfaces in the absence of experimentally determined structures of the IDR/IDP-globular domain complex.

We tested if the AlphaFold2 models of IDRs with high pLDDT scores could be used for rigid-body molecular docking with putative binding partners. To test this, we used as a model system an experimentally determined complex structure of a conditionally folded IDR, the CITED2 transactivation domain (TAD), bound to the folded CBP TAZ1 domain (De Guzman et al. 2004, 2005; Freedman et al. 2003) (PDB: 1p4q), since this enables a comparison to the experimental structure of the complex. Indeed, as we showed above, the AlphaFold2-predicted structure of the CITED2 TAD closely resembles the CBP-bound form (**Figure 3C, 3G, 3K**), with a heavy-atom RMSD between the experimental and AlphaFold2 structures of only 1.6 Å for the entire region and 1.0 Å when aligning the helices only (**Figure 3K**, **Supplementary Figure 4A**). As a control, we first extracted the individual chains for the CITED2 TAD and the CBP TAZ1 in the experimental structure, and rigid-body docked these two chains with protein-protein docking software (**Supplementary Figure 4B**). Reassuringly, the lowest-energy docked structure agreed with the experimental complex (heavy-atom RMSD for residues N216-F259: 0.9 Å), indicating that this strategy could potentially work for AlphaFold2 structures that have atomic-level accuracy to the conditionally folded state of the IDR.

Next, we rigid-body docked the AlphaFold2 structure of the CITED2 TAD onto the experimentally determined structure of the TAZ1 globular domain (**Supplementary Figure 4C**). The results from this simple docking exercise are quite striking: even though the AlphaFold2 structure of the CITED2 TAD closely agrees with the experimental structure of CITED2 (1.6-Å RMSD for all residues or 1-Å RMSD for helices-only), the orientation of the C-terminal helix in the AFDB structure is shifted by 90° relative to the experimental structure (**Supplementary Figure 4A**). The rotation of the C-terminal helix in CITED2 causes a steric clash with the CBP TAZ1 domain that dramatically alters the lowest-energy structure of the docked complex (**Supplementary Figure 4C**), leading to a completely different docked complex. Thus, if one naively used the AlphaFold2 structure of the CITED2 TAD to dock this structure into its interacting globular domain, the resultant molecular model would be seriously flawed.

### Predicted IDRs with high pLDDT scores are enriched in charged and hydrophobic residues

We next sought to understand why AlphaFold2 is folding IDRs/IDPs into high-confidence structures. At least three non-mutually exclusive hypotheses could explain the prevalence of high pLDDT scores in predicted IDRs: (1) global amino-acid sequence differences in comparison to the predicted IDRs with low pLDDT scores, (2) strong signals of co-evolution among residues in these regions, which would imply “high quality” multiple sequence alignments (MSAs) that are unusual for IDRs, and (3) the enrichment of high-pLDDT IDR sequences in the PDB. The first possibility would reflect a differential “folding propensity” that is inherently encoded in the amino-acid sequences of high vs. low pLDDT-scoring IDRs, whereas the latter two possibilities would influence the AlphaFold2 prediction confidence due to the depth of the MSAs (2) or sequence similarity to the structures from the PDB used in training (3) (Jumper et al. 2021a,b). Given the relatively poor coverage of IDRs in the PDB (Quaglia et al. 2021) and the poor positional alignability for most IDRs (Colak et al. 2013; Langstein-Skora et al. 2022; Nguyen Ba et al. 2012; Zarin et al. 2019, 2021), it is plausible that some combination of all three of the aforementioned possibilities could contribute to high pLDDT scoring IDRs.

To gain insight into these possibilities, we first computed the amino-acid frequencies for each of the following three categories: predicted disordered regions with low pLDDT scores below 50 (IDR_low pLDDT_), predicted disordered regions with high pLDDT scores greater than or equal to 70 (IDR_high pLDDT_), and predicted ordered regions (ordered). We hypothesized that the amino-acid frequencies in IDR_low pLDDT_ should reflect the sequence biases found in disordered regions, *i.e.* an enrichment in some charged (D, E, K), polar (Q, S, T), small (G), and helix-disrupting (P) residues (Quaglia et al. 2021). Indeed, the difference between IDR_low pLDDT_ and ordered regions (Δ_ordered_) shows that IDR_low pLDDT_ sequences are enriched in the We next compared IDR_low pLDDT_ and IDR_high pLDDT_ regions (Δ_IDR_) to determine if there are global differences in the sequences of these IDRs that are encoded within per-residue pLDDT scores. Surprisingly, we found that IDR_high pLDDT_ sequences are significantly enriched in E, K, Q, and R residues relative to IDR_low pLDDT_ sequences, as evidenced by the large differences in values of Δ_ordered_ and Δ_IDR_ for these residues (**Figure 6A**). Furthermore, IDR_high pLDDT_ sequences have relatively fewer of some canonical disorder-promoting residues (*e.g.*, P, S, T, D, G) and more order-promoting residues (*e.g.*, C, F, I, L, V, W, Y). Nonetheless, IDR_high pLDDT_ sequences still resemble IDR sequences when analyzed by mean net charge and mean hydropathy (**Supplementary Figure 6**), which is a sequence metric that identifies IDRs from ordered regions (Uversky et al. 2000). However, the IDR_high pLDDT_ sequences show a much broader distribution in both the mean hydropathy and mean net charge dimensions than the IDR_low pLDDT_ sequences (**Supplementary Figure 6**). Thus, although the IDR_high pLDDT_ sequences contain more disorder-promoting residues than ordered regions (**Figure 6B**), IDR_high pLDDT_ sequences appear to have a mixture of both order-and disorder-promoting residues.

**Figure 6.**
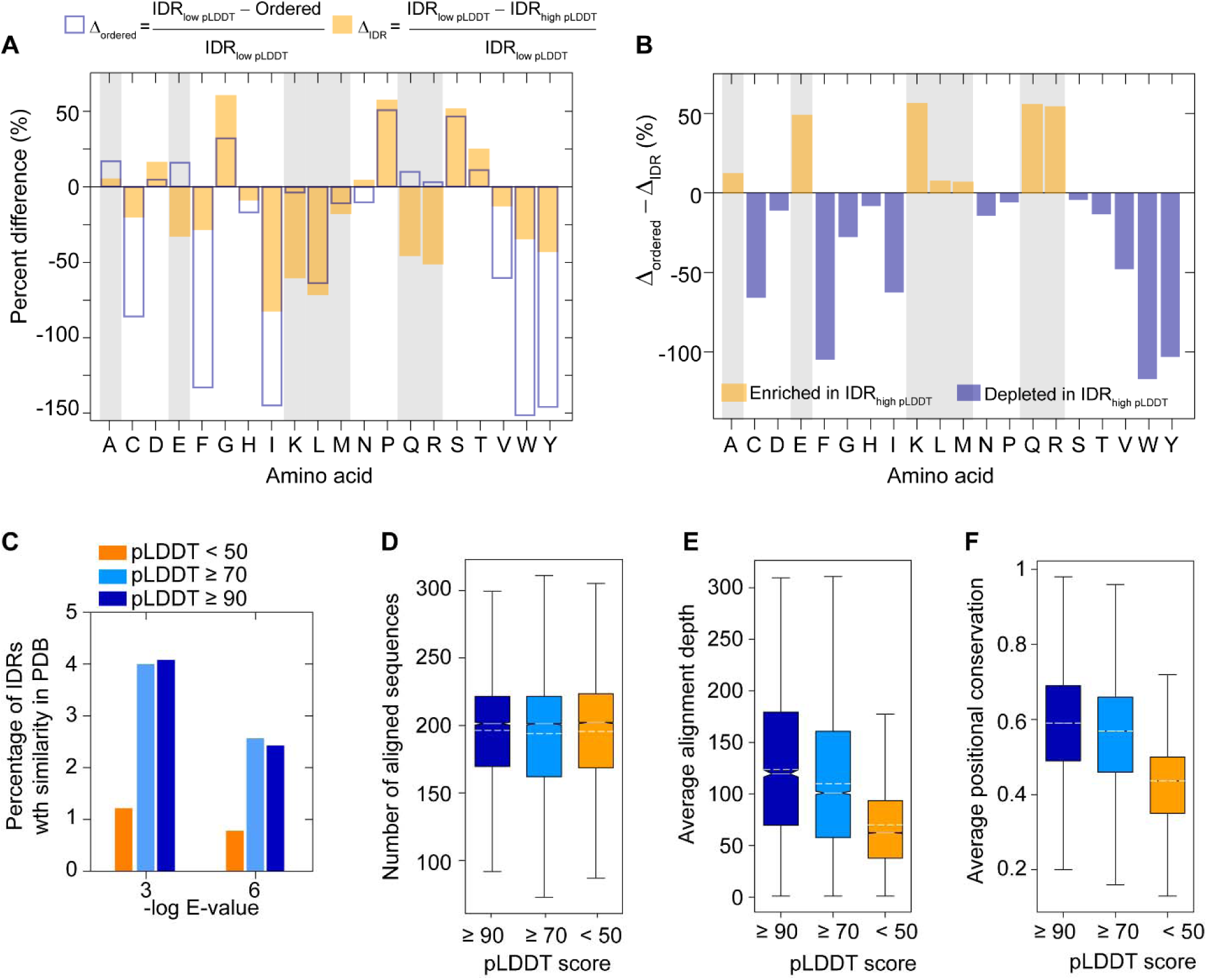
Bioinformatics analysis of predicted IDRs in the AFDB with high pLDDT scores. (**A**) Amino-acid percentages in the regions of predicted order and disorder, with the disordered regions further separated into those with confident pLDDT scores greater than or equal to 70 (IDR_high pLDDT_) and those below 50 (IDR_low pLDDT_). Shown here is the percent change in the relative amino-acid percentages (eq 1) for IDR_low pLDDT_ and either ordered regions (Δ_IDR-order_, empty blue bars) or IDR_high pLDDT_ (Δ_IDR low-high_ orange bars). Positive values indicate that a given amino acid is fractionally enriched in IDR_low pLDDT_ whereas negative values indicate a fractional enrichment in either ordered regions (Δ_IDR-order_) or IDR_high pLDDT_ regions (Δ_IDR low-high_). (**B**) The difference between Δ_IDR-order_ and Δ_IDR low-high_ reports on the relative difference in amino-acid usage between ordered regions and IDR_high pLDDT_ regions as compared to IDR_low pLDDT_ regions. Positive values reflect an increased usage of a given amino acid in IDR_high pLDDT_ regions whereas negative values reflect enrichment in ordered regions with respect to IDR_low pLDDT_ regions, which we assume reflect canonical disordered regions. (**C**) BLASTp results from querying amino-acid sequences in the PDB (**Methods**) for predicted IDRs in the AFDB that are longer than 10 residues. Percentage of predicted IDRs (hits/total) that were identified in the PDB as a function of the *E*-value and the pLDDT score, with < 50 in orange, ≥ 70 in cyan, and ≥ 90 in blue. Box plots of the number of aligned sequences (**D**), average alignment depth (**E**), average positional conservation (**F**). Panel D is not significant whereas E and F have *p*-values (Mann-Whitney) < 0.0001 when comparing pLDDT < 50 and the other groups.

### IDRs with high pLDDT scores have limited similarity to PDB sequences but are more positionally conserved than IDRs with low pLDDT scores

Next, we searched the PDB for IDRs with considerable positional sequence similarity, defined as regions that have over 30% sequence identity over 60% sequence coverage in sequence alignments. We hypothesized that there should be more IDR_high pLDDT_ sequences with similarity to sequences in the PDB than IDR_low pLDDT_ sequences. An enrichment in similarity for IDR_high pLDDT_ sequences could indicate that AlphaFold2 is matching template structures of these IDRs that were used in training. Indeed, we found that IDR_high pLDDT_ sequences are significantly enriched over IDR_low pLDDT_ sequences for similarity to PDB sequences (**Figure 6C**). The percentage of IDRs with pLDDT scores ≥ 70 that have confident BLASTP hits (E-value < 1e-3 or < 1e-6) in the PDB is more than 3-fold higher than IDRs with low pLDDT scores (**Figure 6C**). However, it is important to note that the percentage of high-quality hits in the PDB relative to the total number of predicted IDRs in each pLDDT threshold is very low, *i.e.*, a maximum hit rate of 4% was obtained (**Figure 6C**). Therefore, AlphaFold2 has not simply templated structures of IDRs from the PDB, as the overall coverage of IDR sequences in the PDB remains below 4%.

We next asked if the confident structural predictions for IDRs with high pLDDT scores could reflect higher alignment quality as compared to IDRs with low pLDDT scores. IDRs that conditionally fold have been previously shown to have higher levels of positional amino-acid conservation than IDRs in general (Bellay et al. 2011; Colak et al. 2013). To compute the positional sequence conservation, we constructed MSAs for predicted IDR sequence across different pLDDT categories using homologous sequences retrieved from the ENSEMBL database (Howe et al. 2021). The MSAs contained nearly identical numbers of sequences for each of the three classes of IDRs (pLDDT scores < 50, ≥ 70, and ≥ 90) (**Figure 6D**), yet the average alignment depth was significantly enriched in IDRs with pLDDT scores ≥ 70 and ≥ 90, relative to those with pLDDT scores < 50 (*p*-value < 0.0001) (**Figure 6E**). Moreover, the quality of the alignments was higher for IDRs with high pLDDT scores compared to those with low pLDDT scores, as evidenced by greater levels on average of positional conservation (**Figure 6F**).

Overall, our sequence analysis of predicted IDRs demonstrates that those with high pLDDT scores have higher sequence similarity to the sequences in the PDB than predicted IDRs with low pLDDT scores. However, the overall coverage of IDR sequences in the PDB remains low, with only 4% of high-scoring IDR sequences displaying similarity (E-value < 0.001) to PDB sequences. More significantly, IDRs with high pLDDT scores are more positionally conserved, with nearly 60% sequence identity on average (ignoring gaps; see Methods) and contain fewer gaps than predicted IDRs with low pLDDT scores. Given that AlphaFold2 relies on MSAs as input for its structural predictions (Jumper et al. 2021a), these results provide insight into why AlphaFold2 is folding IDRs with high pLDDT scores into confident structures. The fact that IDRs with high pLDDT scores only rarely have sequence homologs in the PDB suggests that the dominant forces behind the AlphaFold2 predictions for these IDRs are high-quality MSAs and the underlying amino-acid compositions (Toth-Petroczy et al. 2016), and not structural templating.

### AlphaFold2 confidently assigns structures for the majority of IDRs known to conditionally fold

Our bioinformatics analyses provided evidence that IDRs with high pLDDT scores have both compositional differences from and higher quality MSAs than IDRs with low pLDDT scores (Figure 5). IDRs that have high levels of positional conservation are more likely to conditionally fold (Bellay et al. 2011; Colak et al. 2013). Given that our structural analyses above were limited to a handful of conditionally folded IDRs (**Figure 2**, **Figure 3**, **Figure 4**), we sought to gain broader insight into whether the predicted IDRs with high pLDDT scores are, indeed, IDRs that conditionally fold.

To this end, we first investigated the per-residue pLDDT scores for proteins in five databases of conditionally folded IDRs/IDPs across different organisms: the Database of Disordered Binding Sites (DIBS) (Schad et al. 2018), Mutual Folding Induced by Binding (MFIB) (Fichó et al. 2017), DisProt (Quaglia et al. 2021), molecular recognition feature (MoRF) (Disfani et al. 2012), and FuzDB (Hatos et al. 2022) databases. We filtered these databases for regions of IDRs that mapped to the AFDB (Methods) and were left with a total of *ca.* 61,000 residues for further analysis. Remarkably, AlphaFold2 assigned confident pLDDT scores (≥70) to 58.9% of all IDR residues in these databases, ranging from 35% to 87% when each database is analyzed separately (**Supplementary Figure 7**). In comparison, only 14.3% of all residues predicted to be disordered by the SPOT-disorder predictor were assigned confident pLDDT scores (**Figure 1**). Therefore, experimentally validated conditionally folded IDRs are enriched in confident and very confident AlphaFold2 pLDDT scores.

Next, we wondered if the pLDDT scores from AlphaFold2 can be used to differentiate between IDRs that conditionally fold and those that remain disordered. To test this quantitatively, we assessed the classification potential of AlphaFold2 with a receiver operating characteristic (ROC) analysis. We extracted the conditionally folded IDRs from the above-mentioned databases as true positives (*ca.* 61,000 residues). To obtain a dataset of true negatives, *i.e.*, IDRs that do not conditionally fold, we filtered the CheZOD database of proteins with assigned NMR chemical shifts (Dass et al. 2020) to exclude IDRs that have been reported to conditionally fold (see Methods). We were left with *ca.* 10,000 residues in NMR-validated IDRs that are not known to conditionally fold (**Supplementary Figure 8**). We then tested the ability of AlphaFold2 to classify conditionally folded IDRs based on pLDDT scores alone. Using the filtered CheZOD database of IDRs as the true negative dataset, we supplied the five different true positive datasets and found that the ROC analysis yields values of AUC (area under the curve) between 0.58 and 0.90 depending on the input set of true positives (**Figure 7A**, **Supplementary Table 5**). When all of the *ca.* 1400 IDRs that are known to conditionally fold are supplied as input, we find that AlphaFold2 successfully classifies the conditionally folded IDRs with an AUC of 0.72 (**Figure 7A**). Overall, this ROC analysis indicates that the pLDDT scores can, indeed, be used to classify conditionally folded IDRs (**Figure 7A**).

**Figure 7.**
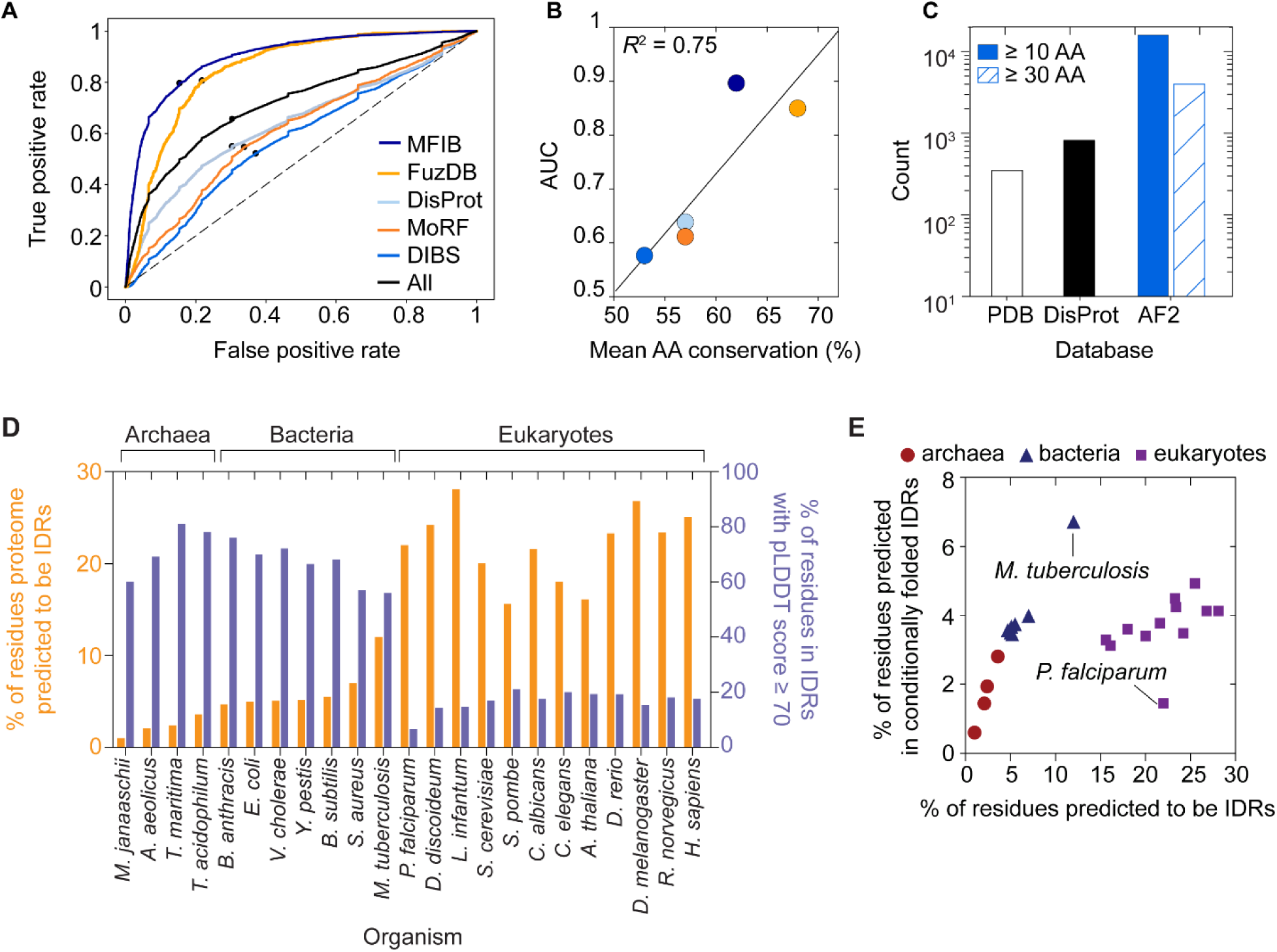
Systematic discovery of conditionally folded IDRs in archaea, bacteria, and eukaryotes. (A) ROC curve for AlphaFold2 pLDDT-based classification of conditionally folded IDRs based on databases of known examples (MFIB, FuzDB, DisProt, MoRF, DIBS). The AlphaFold2-performance on the binary classification task (conditional folder/non-conditional folder) is displayed. Supplementary Table 5 documents performance statistics on all tested databases. All five databases were merged (black) for the classification test on *ca.* 1400 IDRs that are known to conditionally fold. The AUC is 0.72 for the combined dataset. (**B**) Correlation between the mean positional amino-acid conservation of IDR sequences from the databases listed in panel A and the AUC values from the ROC curves in panel A. The best-fit line is shown in black and has a Pearson’s *R*^2^ of 0.75. Note that gaps in the sequence alignments were ignored for the calculation of positional conservation. (**C**) AlphaFold2 expands the sequence and structural coverage of conditionally folded IDRs. The number of human IDRs that are reported to undergo a disorder-to-order transition upon binding is shown (DisProt), but not all of these IDRs have high-resolution structures available. AlphaFold2 increases the number of putative conditionally folded IDRs over DisProt by a factor of 10 or 3, respectively, for continuous regions of 10 or 30 or more amino acids. IDR sequences with appreciable homology to the PDB are shown for comparison. (**D**) For each species listed, the percentage of disordered residues in the proteome (predicted by IUPred2A) is shown in orange on the left y-axis. The percentage of disordered residues with pLDDT scores ≥ 70 (*i.e.*, conditionally folded IDRs) is shown in blue on the right y-axis. (**E**) The percentage of residues in the proteome of each organism from panel A that are both predicted to be disordered and to conditionally fold (pLDDT scores ≥ 70).

When considering the differences in classification performance among the databases, we noticed that the highest AUC values are obtained for MFIB (0.90) and FuzDB (0.85), suggesting that AlphaFold2 has readily learned to identify the conditionally folding segments within these IDRs (**Figure 7A, 7B**). By contrast, the relatively lower AUC values for DisProt (0.64), MoRF (0.61), and DIBS (0.58) indicate that the conditionally folded IDRs within these databases remain somewhat challenging to classify by pLDDT scores alone (**Figure 7A, 7B**). Given the importance of high-quality MSAs to AlphaFold2’s predictions in IDRs (as described above), we tested whether the differences in the performance of AlphaFold2 on various true positive datasets could also be an indication of a reduced evolutionary constraint for conditional folding in some of the IDRs. Indeed, we observed a correlation between the average positional amino-acid sequence conservation of IDRs in each database and the classification performance (AUC) (**Figure 7B**). In other words, the somewhat lower pLDDT scores for conditionally folded IDRs in the DIBS, DisProt, and MoRF databases may reflect higher conformational plasticity enabled by the reduced constraint on positional conservation. Overall, this ROC analysis further supports our hypothesis that IDRs/IDPs with confident and very confident pLDDT scores are likely to be conditional folders.

Given the ability of AlphaFold2 to classify conditionally folded IDRs, we compared the performance of AlphaFold2 as a predictor of conditional folding to that of a software specifically trained for the task of identifying disorder-to-order transitions in IDRs. The software Anchor2 (Mészáros et al. 2018) assigns a score from 0 to 1 to each residue of a given IDR, which represents the probability of a given residue being a part of the disordered binding region that is often coupled to conditional folding or a disorder-to-order conformational transition (Malhis & Gsponer 2015). We observed that the AlphaFold2 pLDDT scores are able to classify conditionally folded IDRs in a manner that is similar or better than Anchor2 (**Supplementary Figure 9**). Furthermore, the classification potential of AlphaFold pLDDT scores to identify conditionally folded IDRs is additionally useful when deployed on a proteome level. Anchor2 reports that nearly 68% of SPOT-Disorder-predicted IDRs are located in binding regions that can potentially undergo a disorder-to-order transition, whereas AlphaFold2 pLDDT scores yield only 14.3% of conditionally folded IDRs. Anchor2 likely also identifies extended short linear interaction motifs (SLiMs) (Benz et al. 2022; Tompa et al. 2014) and other interaction motifs which do not fold upon binding, and thus may be expected to give a larger percentage, not all of which represent conditional folding. This helps understand the Anchor2 value of 68%, which is inconsistent with current knowledge about IDRs and conditional folding (Quaglia et al. 2021). AlphaFold2, in contrast, appears to have learned to identify IDRs that acquire a conditional fold and has encoded this information in the per-residue pLDDT scores.

### AlphaFold2-enabled discovery of conditionally folding IDRs in other organisms

Based on our finding that the pLDDT score of AlphaFold2 enables identification of conditionally folded IDRs, we sought to use AlphaFold2 to discover more IDRs that conditionally fold. In humans there are *ca.* 800 IDRs that are known to conditionally fold (**Figure 7C**) (Fichó et al. 2017; Quaglia et al. 2021; Schad et al. 2018). Our analyses herein have identified nearly 15,000 human IDRs from *ca.* 7,500 proteins that are greater than 10 residues in length that have high-confidence pLDDT scores, which represents an 18-fold increase in the number of known conditionally folded IDRs in the human proteome (**Figure 7C**). Moreover, we identified that only *ca.* 4% of IDRs with high-confidence AlphaFold2 structures have sequence similarity to sequences in the PDB (excluding NMR structures) (**Figure 6B**, **Figure 7C**). Now, with access to more than 8,000 AlphaFold2 structural predictions of IDRs that are putative conditional folders, AlphaFold2 has significantly expanded the structural coverage of conditionally folding IDRs.

In the human proteome, roughly 30% of residues are predicted to be disordered, of which nearly 15% are predicted by AlphaFold2 to conditionally fold. Given that the percentage of intrinsically disordered residues in the proteome has increased from archaea to bacteria to eukaryotes (Gao et al. 2021), we wondered if the percentage of conditionally folded IDRs has also changed. To this end, we used IUPred2A (Mészáros et al. 2018) to predict the IDRs in other AFDBs that are publicly available, as the IUPred2A software program is *ca.* 100-fold faster than SPOT-Disorder and provides a balance between the calculation speed and accuracy of the prediction (Necci et al. 2021). We first compared the predicted number of disordered residues and conditionally folded IDRs in the human proteome, as obtained by using the IUPred2A and SPOT-Disorder predictions to filter the AFDB. We found that IUPred2A and SPOT-Disorder give comparable results: 32% *vs.* 25% of all residues are predicted to be disordered with 14.3% *vs*. 17.6% of predicted disordered residues having pLDDT scores greater than or equal to 70 for SPOT-Disorder and IUPred2A, respectively (**Supplementary Table 1**, **Supplementary Table 2**). Thus, although IUPred2A underestimates the extent of disorder, the boost in calculation speed provides an attractive approach to perform more high-throughput calculations.

We extracted the IUPred2A-predicted IDRs from the 23 AFDBs shown in **Figure 7D**, including four archaeal, seven bacterial, and 12 eukaryotic organisms. As expected, the percentage of disordered residues in the proteome increased from archaea to bacteria to eukaryotes, with minimum and maximum values of 1.0% and 28.1% obtained for *M. janaaschii* and *L. infantum*, respectively (**Figure 7D**). Interestingly, the percentage of IDRs with high-confidence pLDDT scores showed an inverse relation with the overall disordered content. That is, organisms with fewer predicted IDRs have a higher proportion of IDRs with high-confidence pLDDT scores (**Figure 7D**). This result suggests that conditionally folded IDRs are the dominant type of IDRs in the archaea and bacteria examined here, where the percentage of IDRs with high-confidence pLDDT scores ranges from 56% (*M. tuberculosis*) to 81.1% (*T. maritima*). By contrast, in the eukaryotes analyzed here, conditionally folded IDRs appear to be the minority, with minimum and maximum values of 6.6% (*P. falciparum*) and 21.1% (*S. pombe*) (**Figure 7D**). The inverse relation between the percentage of disordered residues in the proteome and the percentage of IDRs that conditionally fold suggests that an upper bound of *ca.* 5% of residues in the proteome localize to conditionally folded IDRs (**Figure 7E**). Although these results on the percentage of IDRs in various proteomes with high pLDDT scores are empirical and presently without theoretical basis, they suggest that organisms with higher percentages of IDRs, and thus a lower fraction of IDRs that conditionally fold, do not functionally utilize IDRs predominantly in the context of conditional folding. Rather, the majority of IDRs in eukaryotic organisms likely function in the absence of conditional folding.

### Leveraging AlphaFold2 to understand disease-causing mutations in conditionally folding IDRs

Our findings indicate that the pLDDT score of AlphaFold2 can be used in combination with sequence-based disorder predictors to identify conditionally folded IDRs. Given these results, we explored whether we could use AlphaFold2 predictions of conditional folding to obtain insight into disease-causing mutations in IDRs.

We computed the per-residue mutational burden in IDRs for disease-*versus* non-disease-associated mutations as a function of the pLDDT score (**Figure 8A**). To this end, we mapped the non-disease-associated sequence variations from the 1000 Genome Project (1000GP) (Auton et al. 2015) to human IDRs (*n* = 332,844 mutations) and calculated the per-residue substitution rates as a function of the range of pLDDT scores (**Figure 8A**). The sequence variants within the 1000GP dataset should predominantly reflect neutral mutations that have risen to appreciable frequency and are therefore polymorphic within the human population. We hypothesized that IDRs with low pLDDT scores, *i.e.* IDRs predicted not to conditionally fold, would be more tolerant to neutral mutations and therefore have a higher per-residue polymorphism rate than IDRs with high pLDDT scores. Indeed, we observed that IDRs with low-confident AlphaFold2 scores have a significantly higher per-residue polymorphism rate than IDRs with high (≥ 70) or very high (≥ 90) pLDDT scores (Fisher Exact Test, *p* < 0.00001 for both comparisons) (**Figure 8A**). Thus, the absence of structural constrains for regions with low pLDDT scores is also reflected in their tolerance to substitutions and more rapid evolution than IDRs with high pLDDT scores (i.e., conditionally folded IDRs) (Pritišanac et al. 2019).

**Figure 8.**
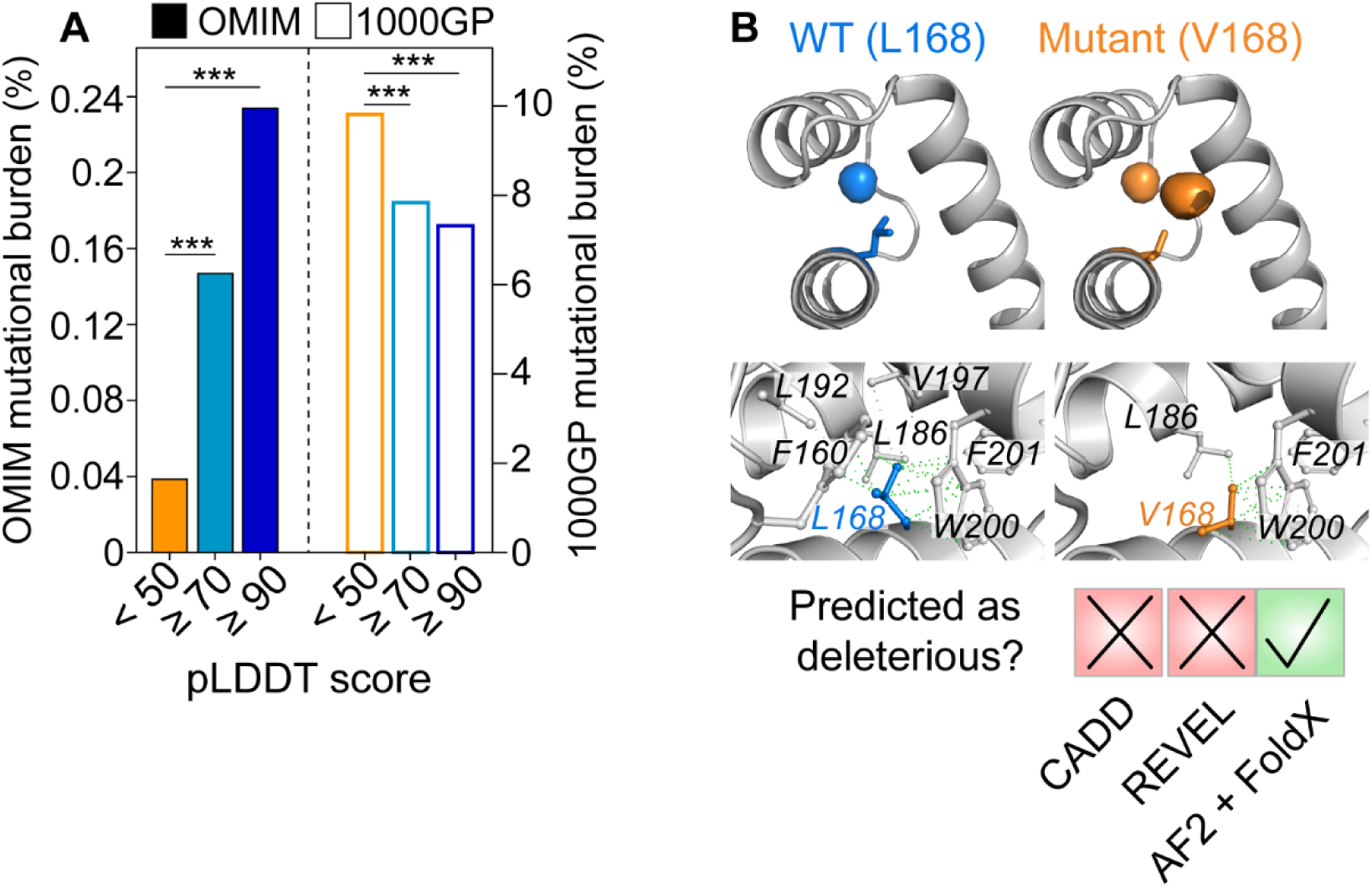
Using AlphaFold2 to understand the basis of disease-causing mutations in conditionally folded IDRs. (**A**) The per-residue mutational burden (the number of mutations divided by the total number of residues) for IDRs is shown as a function of AlphaFold2 pLDDT scores (< 50, ≥ 70, ≥ 90). Disease-associated mutations from OMIM are shown in solid bars, and mutations present in the general human population (1000GP) are shown in empty bars. *** indicates a *p* value < 0.0001 from a Fisher Exact Test. The L168V mutation in *ALX3* causes frontonasal dysplasia, but the mutation is predicted to be “likely benign” by CADD and REVEL. The high-confidence AlphaFold2 structure used in combination with FoldX, however, yields the prediction that L168V is severely destabilizing, with a ΔΔG value of 7.3 ± 0.1 kcal mol^-1^ relative to wild-type ALX3 (ΔΔG = ΔG_WT_ – ΔG_mutant_). Hydrophobic interactions between the L168 side chain (blue) and other atoms in the ALX3 homeodomain that are within a 4.5-Å distance threshold are shown (green lines). A total of 26 interactions were identified. The residues involved in these interactions are indicated. *In silico* mutagenesis of L168 to Val was performed by FoldX. Hydrophobic interactions involving V168 and other atoms in the ALX3 homeodomain are indicated. Only 14 interactions are identified. Arpeggio was used for this analysis (Jubb et al. 2017).

Next, we mapped all known disease-causing mutations from the Online Mendelian Inheritance in Man (OMIM) database (Amberger et al. 2019) to human IDRs (*n* = 1963 mutations). We then calculated the per-residue mutational burden in IDRs that have very high (≥ 90), high (≥ 70), or low (< 50) pLDDT scores (**Figure 8A**). In IDRs with high and very high pLDDT scores, we found a strong enrichment in disease-causing mutations relative to IDRs with low confidence scores (**Figure 8A**). The per-residue mutational burden in IDRs with high and very high pLDDT scores was respectively increased by 4- or 6-fold relative to IDRs with low pLDDT scores (Fisher Exact Test: *p* < 0.00001 for both comparisons). This suggests that, for these disease-associated mutations in OMIM, the IDRs with high confidence AlphaFold2 scores are less tolerant of sequence changes, and that such substitutions are more likely to manifest as disease when compared to IDRs with lower pLDDT scores. Presumably, the restraint of acquiring a conditional fold limits the sequence space of these IDRs and increases the likelihood of a deleterious mutation (Pritišanac et al. 2019).

Moving beyond the statistical association of disease mutations with predicted conditional folding, we next sought to leverage the high-confidence AlphaFold2 structural predictions to gain mechanistic insight into the effects of disease mutations on protein function. An illustrative example is the L168V mutation in the gene *ALX3*, which causes the rare disease frontonasal dysplasia (Twigg et al. 2009). *ALX3* encodes for a transcription factor named homeobox protein aristaless-like 3 (ALX3, UniProt: O95076) that plays a key role in development (Beverdam et al. 2001). The L168V mutation maps to a DNA-binding homeodomain of ALX3 (residues 153-212), for which there is no experimental structure and SPOT-Disorder predicts to be intrinsically disordered. Indeed, the bulk sequence properties of the ALX3 homeodomain, such as mean hydrophobicity (0.37) and mean net charge (0.13), are indicative of an IDR (**Supplementary Figure 10**) (Uversky et al. 2000). Furthermore, other homeodomains are known to be marginally stable in the absence of DNA, *e.g.* the homeodomains from *D. melanogaster* Engrailed and NK-2 have a free energy of denaturation of only 2.2 kcal mol^-1^ (Stollar et al. 2003) and a melting temperature near 25 °C (Tsao et al. 1994), respectively. The binding of DNA to the NK-2 homeodomain induces additional folding and significantly increases its stability (Tóth-Petróczy et al. 2009). Thus, the amino-acid sequence of the ALX3 homeodomain, may encode a conditionally folded IDR.

Even if the ALX3 homeodomain conditionally folds, it remains difficult to rationalize how the chemically subtle L168V mutation might affect function. Indeed, two predictors of variant pathogenicity, the ensemble-based CADD (Combined Annotation Dependent Depletion) (Rentzsch et al. 2019) and the meta-predictor REVEL (Rare Exome Variant Ensemble Learner) (Ioannidis et al. 2016), classify the L168V mutation as “likely benign” (**Figure 8B**). All residues of the ALX3 homeodomain (153-212) have pLDDT scores above 70, and residues 159-208 all have pLDDT scores above 95, indicating a very confident AlphaFold2 prediction with likely accurately positioned side-chain rotamers (Jumper et al. 2021a). Close inspection of the AlphaFold2 structural model reveals that the L168 side chain makes 26 different hydrophobic interactions with nearby aliphatic and aromatic residues from all three helices of the homeodomain (**Supplementary Figure 10**), thus coordinating key inter-helix contacts in the three-dimensional structure. Upon *in silico* mutation of L168 to Val, only 14 of these hydrophobic interactions remain and a new hydrophobic cavity is created (**Figure 8B**).

Since high-confidence AlphaFold2 models perform as well as if not better than experimental X-ray structures for structure-based protein stability calculations (Akdel et al. 2022), we used the high-confidence AlphaFold2 model of the conditionally folded state to calculate protein stability using FoldX (Schymkowitz et al. 2005) (**Figure 8B**). Indeed, the predicted stability of L168V is significantly lower than the wild-type protein, with a FoldX-determined ΔΔG of 7.3 ± 0.3 kcal mol^-1^, nearly 10-fold larger than the error associated with FoldX predictions (**Figure 8B**) (Schymkowitz et al. 2005; Valanciute et al. 2022) and much larger than typical ΔΔG values (1 ± 2 kcal mol^-1^ (Tokuriki et al. 2007)). Thus, the ΔΔG value of 7.3 ± 0.3 kcal mol^-1^ observed for L168V in ALX3, is expected to be highly destabilizing. Given that a positive ΔΔG value indicates an increased population of an unfolded or partially folded state, a likely outcome of the L168V mutation is that the ALX3 homeodomain remains disordered or no longer properly folds upon binding to DNA, thereby preventing or altering its transcriptional activity. Moreover, previous studies have shown that many highly destabilizing mutants cause the protein to be degraded by cellular quality control machinery (Abildgaard et al. 2019; Nielsen et al. 2017). Although the predicted destabilization of ALX3 with the L168V mutation awaits experimental validation, a Leu-to-Val mutation in the homeodomain of a related protein (SHOX) at the same position as L168 in ALX3 was shown to abolish dimerization and DNA binding (Schneider et al. 2005). Our analysis of ALX3 provides a mechanistic hypothesis about how a disease mutation may disrupt the function of ALX3, namely by destabilizing the conditionally folded DNA-bound state.

## Discussion

The application and development of machine learning methods in structural biology has revolutionized the field of protein structure prediction (AlQuraishi 2021; Baek et al. 2021; Evans et al. 2021; Jumper et al. 2021a). AlphaFold2 can predict the structures of most globular proteins to within experimental accuracy (Jumper et al. 2021a). However, approximately 30% of the human proteome is intrinsically disordered, with over 60% of all human proteins containing at least one IDR longer than 30 residues in length (Tsang et al. 2020; Van Der Lee et al. 2014). Thus, it is essential to critically analyze the AlphaFold2-predicted structures of IDRs, as such regions cannot be accurately described by a single, static structure (Ruff & Pappu 2021).

Here, we have shown that there are thousands of predicted IDRs in the human proteome that are ascribed confident or very confident pLDDT scores in the AFDB (**Figure 1A**), and thus have confidently predicted three-dimensional structures. These high-confidence AlphaFold2 structures of IDRs often can capture a folded conformation that forms in the presence of a specific binding partner or upon post-translational modification (**Figure 2**, **Figure 3**, **Figure 4**), even though the AlphaFold2 structures were predicted in the absence of such binding partners or post-translational modifications. AlphaFold2 assigns confident pLDDT scores to these IDRs likely due to constraints imposed by their amino-acid sequences, and not PDB templating, as we find that 96% of the IDR sequences with high confidence AlphaFold2 structures do not have appreciable similarity to the PDB (**Figure 6C**). Thus, AlphaFold2 has likely identified conditionally folded IDRs through the co-evolution of specific residues, which can indicate spatial proximity within a three-dimensional structure (Anishchenko et al. 2017; De Juan et al. 2013; Göbel et al. 1994; Karamanos 2023; Morcos et al. 2011). Indeed, co-evolution can recover the folded structures of some conditionally folding IDRs (Tian et al. 2015; Toth-Petroczy et al. 2016).

Expanding our analysis of conditionally folded IDRs, we further found that AlphaFold2 can classify conditionally folded IDRs based solely on the per-residue pLDDT score. The performance of AlphaFold2 as a classifier of conditional folding depended on the input IDRs, with those in the MFIB (AUC = 0.90) and the DIBS (0.58) as the extrema. All databases were assembled through manual curation of the literature and publicly available databases, and not through sequence similarity. Thus, one possible explanation for the difference in classification performance is that the conditionally folded IDRs that AlphaFold2 struggles to identify might be more conformationally dynamic or sample more than one structure in the conditionally folded state. If so, there may be less of an evolutionary constraint for stable structure, which may lower the final pLDDT scores assigned to such regions.

A valuable metric that emerges from our analyses is an estimate on the fraction of IDR residues that conditionally fold (**Figure 7C**). There are presently *ca.* 800 known human IDRs that conditionally fold, based on manual curation of the DisProt database. Thus, the *ca.* 15,000 human IDRs longer than 10 residues with high-confidence AlphaFold2 structures provide a useful resource to interrogate the sequence and structural properties of IDRs that acquire folds during their function. At present, we do not know the rate with which AlphaFold2 incorrectly predicts conditionally folded IDRs; however, noting that AlphaFold2 correctly identified *ca.* 60% of IDRs that are known to conditionally fold over five different databases, we can estimate an upper bound of conditionally folded human IDRs at 25% (0.15/0.60). A percentage of conditionally folded human IDRs between 15-25% correlates with previous observations that a minority of human IDRs display significant degrees of positional sequence conservation (Colak et al. 2013), with many of the positionally conserved IDRs identified as those that fold upon binding (Bellay et al. 2011; Colak et al. 2013). Positional conservation is atypical of IDRs that generally evolve rapidly and show low positional conservation, even though there is significant conservation of bulk molecular features (Zarin et al. 2019, 2021). Therefore, there may be purifying selection upon IDR sequences that function by conditional folding, such that the sequence only slowly evolves in order to maintain the overall fold of the bound/modified form of the IDR.

Collectively, these results lead to the hypothesis that if only 15-25% of human IDRs are conditionally folded, then the majority of IDRs in the human proteome, and those of other eukaryotes, would function in the absence of stable structure. These would include IDRs involved in discrete dynamic or “fuzzy” complexes (Borgia et al. 2018; Fuxreiter 2019; Mittag et al. 2008; Tompa & Fuxreiter 2008) and those with low complexity sequences that participate in dynamic, exchanging condensed phases of biomolecular condensates (Murthy & Fawzi 2020; Peran & Mittag 2020). On the other hand, our analysis of the percentage of conditionally folded IDRs in various organisms reveals that archaea and bacteria, which have lower disordered content throughout the proteome (Gao et al. 2021), have a relatively higher percentage of IDRs that conditionally fold (**Figure 7F, 7G**). In prokaryotes, therefore, the majority of IDRs seem to conditionally fold.

Rather than a single AlphaFold2 model, a more suitable structural representation of IDRs requires conformational ensembles that describe the free energy landscapes of IDRs, and thus their accessible conformations (Davey 2019; Jensen et al. 2014; Mittag & Forman-Kay 2007; Pietrek et al. 2023; Wei et al. 2016). The structural plasticity of IDRs and the interconversion between different structural states is essential to their biological function (**Figure 3**). If AlphaFold2 can recover the heterogeneity in conditionally folded IDRs fold into multiple conformations, such as p53 (Berlow et al. 2015), then these predictions would be valuable to search for plausible structured conformations that are accessible to a given IDR. Interestingly, recent studies found that reducing the depth of the MSA, performing rational *in silico* mutagenesis within the MSA, or clustering the MSA can drive AlphaFold2 to sample alternate conformations (Alamo et al. 2021; Stein & Mchaourab 2021; Wayment-Steele et al. 2022). However, it remains to be tested if this approach can recover the structural landscape of conditionally folded IDRs that fold into different conformations.

The development of small molecules that target the binding sites for IDRs, as well as the IDRs themselves, is an active area of pharmaceutical research (Metallo 2010). Because IDRs/IDPs are enriched in many signaling pathways (Wright & Dyson 2015), they are highly desirable drug targets (Metallo 2010); however, it is notoriously difficult to target IDRs/IDPs with pharmaceuticals, which likely is due to their absence of stable structures, large interaction networks (Teilum et al. 2021), and interconversion between different structural forms (Metallo 2010). Having access to high-confidence AlphaFold2-predicted structures for the *ca.* 15% of conditionally folded IDRs/IDPs may accelerate the targeting of IDRs/IDPs with rationally designed inhibitors.

Our finding that AlphaFold2 has learned to identify conditionally folded IDRs will enable new insights into the sequences, evolution, and structural bioinformatics of these regions. Along with other recent analyses of IDRs (Brotzakis et al. 2023; Piovesan et al. 2022; Ruff & Pappu 2021; Wilson et al. 2022), our work demonstrates that AlphaFold2 has learned interesting properties about certain IDRs.

While the structural predictions generated by AlphaFold2 will certainly accelerate the pace of biomedical discovery, there remains a huge need for experimental (Bhowmick et al. 2016) and bioinformatic (Zarin et al. 2019, 2021) approaches to address the majority of IDRs that likely function in the absence of folded structure. With increased experimental data on IDRs/IDPs, including integrative structural modelling (Bottaro et al. 2020; Choy & Forman-Kay 2001; Gomes et al. 2020; Krzeminski et al. 2013; Lincoff et al. 2020; Ozenne et al. 2012; Salmon et al. 2010), machine-learning methods promise to provide new insights into disordered protein conformational states and functional mechanisms (Lindorff-Larsen & Kragelund 2021). The complementary nature of AlphaFold2 structural predictions and atomic-level insight from NMR spectroscopy will be pivotal for future developments related to IDR conformational ensembles.

## Methods

### Sequence-based prediction of IDRs in the human proteome

We obtained all protein sequences from the human proteome from the UniProt database (reference proteome number UP000005640, downloaded in November 2021). This reference human proteome contains 20,959 unique UniProt IDs that have a total of 11,472,924 residues. To identify IDRs in the human proteome, we used SPOT-Disorder (Hanson et al. 2017), which was recently identified as one of the most accurate predictors of disorder (Necci et al. 2021) and gave the closest agreement with experimentally determined intrinsic disorder based on NMR data (Dass et al. 2020; Nielsen & Mulder 2019). Regions of the proteome that were not predicted to be disordered were assumed to be ordered. For analysis of the per-residue pLDDT scores, the SPOT-Disorder predictions were used without filtering for consecutive residue length (**Figure 1A**); in the bioinformatic analyses of **Figure 6**, to exclude very short segments, we filtered the SPOT-Disorder predictions to include only the regions with predicted consecutive disorder greater than 10 residues.

For the analysis of sequence-based predictors of disorder in **Figure 2A**, we used the software packages metapredict, SPOT-Disorder, DISOPRED3, and IUPred2A (Emenecker et al. 2021; Hanson et al. 2017; Jones & Cozzetto 2015; Mészáros et al. 2018). SPOT-Disorder was run as noted above. The webserver versions of the other three software programs were used with default parameters.

### Extraction of per-residue pLDDT scores from the AFDB

Per-residue pLDDT scores were extracted from each PDB file in the database using an in-house Python script. The per-residue SPOT-Disorder predictions of disorder were then mapped onto the AFDB in order to split the AlphaFold2 data into predicted regions of disorder and order. In this way, our analysis should not be biased by the assignment of secondary structure in the AlphaFold2-generated structures; instead, we relied on SPOT-Disorder to identify the IDRs and ordered regions. The SPOT-Disorder-predicted regions of disorder were further split into the AlphaFold2-designated levels of confidence: very low (≤ 50), low (≤ 70), confident (≥ 70 *x* < 90), or very confident (≥ 90) pLDDT scores.

For the organisms listed in **Figure 7**, AFDBs were downloaded from the AlphaFold website in January 2022. For organisms that did not have pre-compiled AFDBs at that point, but did have AlphaFold2 structures available, an in-house Python script was used to automatically download all structures that matched query UniProt IDs from a UniProt proteome file.

### PDB files within the AFDB

The repository of human protein structures was downloaded in November 2021 from the AlphaFold Protein Structure Database using the reference proteome number UP000005640. This corresponded to version 1 of the human AFDB, as indicated with the “v1” string in the name of the downloaded file. The database contained 23,391 predicted structures that map to 20,504 unique UniProt IDs, which corresponds to 97.8% of the proteome (containing 20,959 unique entries). The total number of residues in the AFDB is 10,825,508, which is 94.4% of all residues in the proteome (11,472,924 residues) and *ca.* 2.6% more than what was reported in the original work (Tunyasuvunakool et al. 2021), likely reflecting the *ca.* 1% increase in the number of UniProt IDs (20,504 in AFDB *vs.* 20,296 sequences in the original work).

In the AFDB, the greater number of structures (23,391) relative to the number of unique UniProt IDs (20,504) is due to the 2,700-residue limit for AlphaFold2 structures: proteins longer than this threshold are segmented into multiple structures that contain overlapping 1,400-residue fragments (*e.g.*, residues 1-1400, 201-1600, etc.). The human proteome contains 210 proteins longer than 2,700 residues; searching the AFDB for the UniProt IDs that map to these proteins reveals 3,095 AlphaFold2 structures, accounting for the difference in the number of PDB files and unique UniProt IDs.

### Biological Magnetic Resonance Bank

Assigned NMR chemical shifts for the IDRs/IDPs α-synuclein, 4E-BP2, and ACTR were downloaded from the BMRB (Ulrich et al. 2008) using the following entry identification numbers: 6968 (α-synuclein) (Bermel et al. 2006), 5744 (α-synuclein bound to SDS micelles) (Chandra et al. 2003), 19114 (4E-BP2) (Lukhele et al. 2013), 19905 (phosphorylated 4E-BP2) (Bah et al. 2015), 15397 (ACTR) (Ebert et al. 2008), and 5228 (ACTR bound to CBP) (Demarest et al. 2002). All of the entries contained assignments for ^1^HN, ^1^Hα, ^15^N, ^13^CO, ^13^Cα, and ^13^Cβ chemical shifts, except for 5744 (no ^1^Hα and ^13^CO assignments) and 15397 (no ^1^Hα assignments).

### Calculation of NMR chemical shifts from an input PDB structure

To simulate the NMR chemical shifts of AlphaFold2-generated structure predictions, we used the SPARTA+ software package (Shen & Bax 2010). Protons were first added to each PDB structure using DYNAMO version 7.2 available via the PDB Utility Web Servers from the Bax Laboratory. The proton-containing PDB files were then uploaded to the SPARTA+ Web Server using default parameters. The backbone and ^13^C^β^ chemical shifts were extracted from the output file with an in-house Python script.

### Calculation of secondary structure propensity from NMR chemical shifts

We used the SSP software program (Marsh et al. 2006) via the NMRbox (Maciejewski et al. 2017) to calculate the secondary structure propensity (SSP) of the IDRs/IDPs shown in **Figure 2**. We included ^13^C^α^ and ^13^C^β^ shifts as input, as recommended in the original SSP publication. Prior to analysis, we re-referenced all of the chemical shifts using standard protocols (Marsh et al. 2006). This is important because secondary chemical shifts, and therefore SSP, are highly sensitive to the internal referencing of the measured chemical shifts, and any errors in referencing will impact the downstream SSP analysis. The SSP-derived re-referencing offset ppm in the ^13^C dimension for each dataset was -0.390 (BMRB: 6968), -0.483 (5744), +0.373 (191141), +0.106 (19905), -0.051 (15397), and -0.139 (5228).

### Sequence similarity between IDRs and the PDB

We ran BLASTP (Altschul et al. 1990) to determine to which extent IDR sequences with different pLDDT scores overlap with sequences in the PDB. To filter the sequences that are in the PDB, we removed duplicates of identical sequences and highly homologues sequences using PISCES (Wang & Dunbrack 2003) with the following parameters: maximum pairwise percent sequence identity (75%), resolution (0-3.5A), minimum chain length (40), and the maximum chain length (10,000). The sequence data included all X-ray structures that met the aforementioned criteria but excluded all NMR and cryo-EM entries as well as sequences that map to regions of X-ray structures with no electron density. We separately downloaded all X-ray structures that contain 40 or fewer residues. The curation resulted in a set of *ca.* 4,000 unique sequence entries, which were then used to create the BLAST database. BLASTP was run with E-value cut-off values of 1e-3 and 1e-6, with restrictions on sequence identity (>30%) and coverage criteria (>60%).

### Evaluation of positional sequence conservation in IDRs

Positional sequence conservation was computed for alignments of IDRs that were distributed in three sets with different cut-offs of pLDDT scores (see “Mapping pLDDT scores to IDRs”). Only IDR sequences with 10 or more residues with consecutive pLDDT score below or above the desired threshold were considered. To compute positional conservation across MSA columns, we used a modified metric of Shannon’s entropy, the so-called property entropy as introduced by Capra and Singh (Capra & Singh 2007). Gaps were ignored in the computation of positional conservation.

### Bioinformatic analysis of the predicted IDRs in the AFDB

There are 10,825,508 residues in the AFDB, of which 3,539,799 are predicted by SPOT-Disorder to be disordered (**Supplementary Table 1**), 7,127,685 to be ordered, and 158,024 are not mapped. The latter is consistent with the *ca.* 98.5% coverage of the human proteome in the AFDB (Tunyasuvunakool et al. 2021). From these numbers, we can also calculate the percentage of residues in SPOT-Disorder-predicted IDRs, which amounts to 32.7% of the AFDB, in agreement with literature values (Necci et al. 2021).

Next, the SPOT-Disorder-predicted IDRs were further split into those with low pLDDT scores (< 70) and those with high pLDDT scores (≥ 70). A total of 506,101 residues are in IDR_high pLDDT_ (pLDDT ≥ 70) as compared to 3,033,698 in IDR_low pLDDT_ (pLDDT < 70). Using an in-house Python script, we calculated the amino-acid frequencies in each of the following categories: ordered regions, IDRs with low pLDDT scores (IDR_low pLDDT_), and IDRs with high pLDDT scores (IDR_high pLDDT_). The amino-acid frequencies were normalized to the total number of amino acids in each category to yield the percentage of the total:

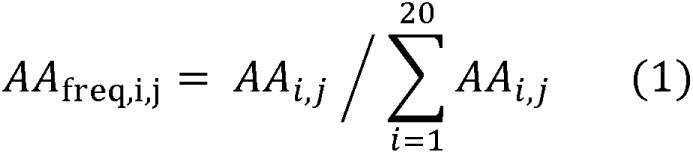

Where *AA*_i,j_ stands for the number of amino acids of residue type *i*, with the indices *i* and *j* respectively indicating the amino acid type (A, C, D, …, V, W, Y) and the category of residues analyzed (ordered, IDR_low pLDDT_, IDR_high pLDDT_). The summation in the denominator of eq 1 refers to the total number of amino acids in each category.

Mean net charge and mean hydrophobicity of IDR_low pLDDT_ and IDR_high pLDDT_ sequences were calculated according to Uversky *et al*. (Uversky et al. 2000) using an in-house Python script. Briefly, the net charge of a given IDR/IDP sequence was computed at pH 7. The p*K*_a_ of histidine residues was set to 6.5 to match an experimental determination of the His p*K*_a_ in an IDP recorded in the presence of physiological salt concentrations (Croke et al. 2011). The absolute value of the net charge of the IDR/IDP was then divided by the total number of residues to obtain the mean net charge. The mean hydrophobicity of an IDR was computed using a normalized version of the Kyte-Doolittle hydropathy scale, such that the values ranged between 0 and 1. The hydropathy value of each residue in the IDR/IDP was then averaged over a sliding window of five residues. The mean hydropathy was finally obtained by computing the sum of all hydropathy values (via the sliding window) and then dividing by the total number of residues.

### Databases of IDRs/IDPs that fold upon binding

To determine if AlphaFold2 can systematically identify IDRs/IDPs that conditionally fold, we first extracted the amino-acid sequences of IDRs/IDPs that are known to conditionally fold (i.e., true positives). To this end, we filtered five databases that contain manually curated lists of IDRs/IDPs that conditionally fold: DisProt (Quaglia et al 2022), FuzDB (Hatos et al 2021), and the previously mentioned DIBS (Schad et al 2018) and MFIB (Fichó et al 2017). Then, in order to extract the per-residue pLDDT scores of these IDR/IDP sequences, an in-house Python script was written to download and filter AlphaFold2 structural models from the AFDB. From the DIBS and MFIB databases 551 and 253 regions accounted for 9,351 and 20,482 residues, respectively. Within the DisProt database, we only considered IDRs that have the annotation “structural transition: disorder-to-order”, which generated a total of 473 regions and 27,013 residues with corresponding pLDDT scores for further analysis. Finally, from 95 regions in the FuzDB that are listed as “disorder-to-order regions” (DOR), 757 pLDDT scores of corresponding residues were extracted for further analysis. The output files contained the UniProt IDs, pLDDT scores, amino-acid types, and residue numbers of the IDRs/IDPs that were taken from each database.

Next, we compiled a curated dataset of IDRs/IDPs that do not fold upon binding (i.e., true negatives). Previous software programs that were specifically designed to detect disordered binding regions were trained on a list of flexible linkers between structured domains (Disfani et al. 2012; Dosztányi et al. 2009; Mészáros et al. 2018). From this file, 386 regions were obtained with a total of 4,765 residues. However, when we examined the AlphaFold2 pLDDT scores of these flexible linkers, we found that the majority of these regions are short (e.g., fewer than 10 residues) and highly conserved with high or very high pLDDT scores (**Supplementary Figure 8**) For these reasons, we constructed a new dataset of true negatives using NMR data from IDRs that have not been reported to conditionally fold. We used the CheZOD database (Nielsen & Mulder 2016) to identify IDRs/IDPs that do not conditionally fold. Importantly, CheZOD contains a manually curated and filtered list of proteins with assigned NMR chemical shifts that are used to experimentally quantify (dis)order at the residue level (Nielsen & Mulder 2020).

The expanded version of the CheZOD database contains experimental NMR data from 1325 protein sequences (Dass et al. 2020). Two files were downloaded from the CheZOD database: the first one matched the BMRB ID with the corresponding amino-acid sequence, whereas the second file contained only the raw Z-scores without the corresponding BMRB ID or amino-acid sequence. Thus, to relate the sequences with their Z-scores and UniProt ID, we used BLASTp to query each sequence, from which we retrieved the corresponding UniProt ID. The resulting file matched each sequence with the most confident BLASTp hits (E-value < 1e-3 or <1e-6) and contained information about the UniProt ID, the range of amino acids in the queried sequence, which was aligned to the reference sequence, and the matching amino-acid range in the aligned sequence. Since CheZOD was trained on both ordered and disordered sequences, we first removed regions with secondary or tertiary structure (Z-score > 3.0). We filtered this set of sequences to retain only those residues that have Z-scores below 3.0, which is indicative of disorder, as recommended by the developers (Dass et al. 2020). To extract only the unstructured regions, we then matched the remaining Z-scores (< 3.0) and residues with their UniProt IDs. The output file contained the UniProt ID and amino acid range of disordered regions. However, some UniProt IDs contained regions shorter than 5 consecutive residues, which we then also removed. Finally, given that CheZOD contains protein sequences with associated NMR data, and thus is agnostic to the conditional folding of IDRs/IDPs, we excluded from CheZOD any sequence that overlaps with any of the five databases of IDRs/IDPs that conditionally fold. The final, filtered CheZOD database contained 557 regions. After projecting those regions to their associated AlphaFold2 structural models, a total of 9,534 pLDDT scores were extracted. In this way, by filtering CheZOD to only retain sequences with NMR-derived evidence of disorder, and by then removing any known conditionally folded IDRs/IDPs, we compiled a list of IDRs/IDPs that are experimentally validated and have not been reported to conditionally fold. Despite our stringent filtering, some sequences within this dataset may conditionally fold (false negatives), but this cannot be avoided. We found that AlphaFold2 assigns low-confidence pLDDT scores (< 70) for nearly 80% of the residues in our filtered CheZOD database, which resembles the proportion of IDRs in human proteome that are given low-confidence pLDDT scores (∼85.7%, **Figure 1B**). Moreover, the IDRs within the filtered CheZOD database have a longer average length and a lower average alignment depth than IDRs in the flexible linkers dataset (**Supplementary Figure 8**).

Finally, we used these true negative (filtered CheZOD) and true positive databases (MFIB, FuzDB, DIBS, DisProt, MoRF) to assess the performance of AlphaFold2 on identifying IDRs/IDPs that conditionally fold. We used the pLDDT scores of the extracted regions that conditionally fold from each database as a true positive dataset, whereas the pLDDT scores of regions extracted from CheZOD database that do not conditionally fold were used as a true negative dataset. The expected pLDDT scores were all set to 1 for the true positive dataset and to 0 for the true negative dataset. We then plotted ROC curves by comparing the observed pLDDT scores (normalized between 0 and 1) from the AlphaFold2 structural predictions against the expected pLDDT scores (0 or 1). The associated AUC, precision, and recall values for each true positive database are listed in **Supplementary Table 5**.

We ran the Anchor2 software with the default parameters on all sequences in the aforementioned databases (Mészáros et al. 2018). Anchor2 was originally developed to identify regions in IDRs that bind to other proteins, including those that undergo a disorder-to-order transition. Although not specifically designed to detect conditional folding, we used Anchor2 to compare to AlphaFold2 for the purpose of identifying IDRs that fold in the presence of a binding partner or upon PTM. We then performed an ROC analysis as above in which the values for IDRs in the same true positive and negative datasets were set to 1 and 0, respectively. Except for the DIBS database (AUC 0.59, precision 0.57, recall 0.61), which was used in the training of Anchor2, the classification of conditionally folded IDRs remains a challenging task for Anchor2. Overall, our classification comparison shows that, even though AlphaFold2 was never specifically trained to detect conditional folding or possible binding sites within IDRs, it performs well on this task, regardless of the differences in types of IDRs from different databases (**Supplementary Figure 9**).

### Simulation of biophysical parameters from an input PDB structure

As outlined in **Figure 5**, **Supplementary Figure 3,** and the **Supplementary Appendix**, biophysical experiments can be performed and compared to predictions derived from the AFDB structural model. Such comparisons would be able to rapidly report on the accuracy of the model relative to the conformations sampled in solution.

#### Circular dichroism (CD)

CD spectra of proteins are sensitive to global secondary structure content and do not require much sample. The webserver PDB2CD (Mavridis & Janes 2017) was used with default parameters to simulate the CD spectrum of the AFDB structure of human α-synuclein and to compare to experimental data (Fusco et al. 2016).

#### Translational diffusion (PFG-NMR)

The software package HYDROPRO was used to calculate hydrodynamic properties of α-synuclein based on its structure in the AFDB (Ortega et al. 2011) (**Supplementary Figure 3B**). Unless otherwise specified, default parameters were used (*e.g.,* a non-overlapping shell model was used with shell model set to the atomic level and a 2.84-Å radius of atomic elements). The temperature was set to 15 °C to match the experimental conditions (Ramanujam et al. 2020b), and the solvent viscosity was adjusted accordingly to 1.1366 cP based on the value reported by NIST. The AlphaFold2 structure from the AFDB was then loaded and the calculation was run. The predicted translational diffusion coefficient from HYDROPRO is 5.141 x 10^-11^ m^2^ s^-1^, which is approximately 10% smaller than the experimentally measured value of 5.71 ± 0.02 x 10^-11^ m^2^ s^-1^ (Ramanujam et al. 2020b), suggesting that the AlphaFold2 structure is more extended than the conformation of α-synuclein observed experimentally. To generate a simulated plot of signal decay *(I*_j_ / *I*_0_) caused by translational diffusion during a BPP-LED pulse sequence, equation 2 below was used:

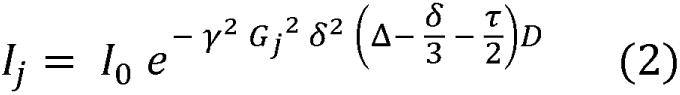

Where the measured signal intensity, *I*_j_, depends on the exponential term above that contains the square strength of the applied gradient (*G*_j_^2^), which is varied during the experiment, and a linear contribution from the translational diffusion coefficient (*D*). The other parameters are either held fixed in the experiment (Δ, the delay time for translational diffusion; δ, the total duration of the encoding gradients; τ, the gradient recovery duration) or are physical parameters (γ, the gyromagnetic ratio of ^1^H). Values of 267,522,187.44 rad s^-1^ T^-1^, 200 ms, 3 ms, 200 μs, and 0.668 T m^-1^ were used for γ, Δ, δ, τ, and *G*_max_, respectively, to match experimental conditions (Ramanujam et al. 2020b). The values of *G*_j_ were varied over a range of *G*_j_ / *G*_max_ from 0 to 1.

#### Small-angle X-ray scattering (SAXS)

The software program Crysol (Svergun et al. 1995) was used to simulate SAXS data of human α-synuclein based on the AFDB structure (**Supplementary Figure 3C**). Experimental SAXS data are available for comparison (Ahmed et al. 2021). The software package ATSAS (Manalastas-Cantos et al. 2021) was used to analyze the SAXS data to obtain the radius of gyration (*R*_g_) and the maximum distance (D_max_). The fitted experimental values of *R*_g_ and *D*_max_ are 35.6 ± 0.2 Å and 109 Å, and those derived from the simulated data from the AFDB structure are 42.6 Å and 152 Å, respectively.

#### Other NMR parameters

The ^13^Cα chemical shifts in **Supplementary Figure 3D** were simulated with SPARTA+ (Shen & Bax 2010) after protons had been added to the AFDB structure of human α-synuclein and neighbor-corrected random coil chemical shifts were obtained from the SPARTA+ output file. The measured ^13^Cα chemical shifts were extracted from (Bermel et al. 2006). The ^3^*J*_HNHα_ coupling constants in **Supplementary Figure 3E** were simulated from the AFDB structure of human α-synuclein in which protons had been added, as described above. The parameterized form of the Karplus equation used to relate the dihedral angle Φ to the ^3^*J*_HNH_ coupling constant is shown below and based on (Vögeli et al. 2007):

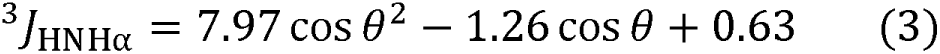

Where B is the dihedral angle Φ minus 60°, which is then converted to radians. Dihedral angles were computed from the AFDB structure of human α-synuclein using an in-house Python script that uses the BioPython package (Cock et al. 2009). The simulated values of ^3^*J*_HNH_ were compared to those measured experimentally (Mantsyzov et al. 2015). Finally, ^1^Hα solvent paramagnetic relaxation enhancements (sPREs) were obtained from (Hartlmüller et al. 2019). Simulated sPREs were performed using the sPRE-calc software program (Gong et al. 2017), and the simulated rates were then scaled to roughly match the lowest experimentally reported sPRE values.

### Disease mutations

The missense variants from the OMIM database (Amberger et al. 2019) annotated in UniProtKB were available for download on the UniProt website (humsavar.txt, September 2019). An in-house Python script was used to map the OMIM missense mutations to SPOT-Disorder predicted IDRs that are greater than or equal to 10 residues in length. We next assessed the differences in per-residue missense mutation rates in IDRs with different pLDDT scores, as annotated in the AFDB. Specifically, we focused on the difference in the per-residue mutation rates in IDRs with very high (≥ 90), high (≥ 70), or low (< 50) pLDDT scores. As a control, we also assessed the per-residue mutation rates of neutral variants from the 1000 Genome Project (1000GP) (Auton et al. 2015), which were annotated in UniProtKB and available for download from the UniProt website (homo_sapiens_variation.txt, September 2019). As above, we mapped the missense variants to IDRs that were greater than or equal to 10 residues in length, and we then split the IDRs into three categories based on pLDDT scores. The Fisher Exact Test was used to assess the significance of the difference in per-residue mutation rates in regions with very high and high pLDDT scores as compared to regions with low pLDDT scores. In all four comparisons (OMIM: very high vs. low, high vs. low; 1000GP: very high vs. low, high vs. low), the *p* value was less than 0.0001.

### FoldX

The AlphaFold2 structural model of ALX3 was downloaded from the AFDB. Residues corresponding to the homeodomain (152-213) were excised and saved as a separate PDB file. The homeodomain PDB file was then supplied as input for the FoldX command “RepairPDB”. Version 4 of FoldX was used (Schymkowitz et al. 2005). The mutation L168V was introduced into the output PDB file by the FoldX command “BuildModel”, with the number of runs set to five. The average ΔΔG value and standard deviation were computed over these five runs. The FoldX protocol was based on (Valanciute et al. 2022). Inter-atomic interactions in the wild-type and mutant PDB files were analyzed with the Arpeggio webserver (Jubb et al. 2017). Cavity detection was performed in PyMol with a cavity detection radius and cavity detection cutoff both set to 3 solvent radii.

## Data Availability

AlphaFold Protein Structure Databases (AFDBs) were downloaded from the AlphaFold EMBL-EBI website (https://alphafold.ebi.ac.uk/). UniProt Proteome files were downloaded from UniProt (https://www.uniprot.org/) for *Aquifex aeolicus* (UP000000798), *Arabadopsis thaliana* (UP000006548), *Bacillus anthracis* (UP000000594), *Bacillus subtilis* (UP000001570), *Candida albicans* (UP000000559), *Caenorhabditis elegans* (UP000001940), *Dictyostelium discoideum* (UP000002195), *Drosophila melanogaster* (UP000000803), *Danio rerio* (UP000000437), *Escherichia coli* (UP000000625), *Homo sapiens* (UP000005640), *Leishmania infantum* (UP000008153), *Methanocaldococcus jannaschii* (UP000000805), *Mycobacterium tuberculosis* (UP000001584), *Neurospora crassa* (UP000001805), *Plasmodium falciparum* (UP000001450), *Rattus norvegicus* (UP000002494), *Saccharomyces cerevisiae* (UP000002311), *Schizosaccharomyces pombe* (UP000002485), *Staphylococcus aureus* (UP000008816), *Synechocystis sp.* (UP000001425), *Thermoplasma acidophilum* (UP000001024), *Thermotoga maritima* (UP000008183), *Vibrio cholerae* (UP000000584), and *Yersinia pestis*(UP000000815).

The specific UniProt IDs for the proteins discussed in the text are: ataxin-8 (Q156A1), complexin-3 (Q8WVH0), synaptosomal-associated protein 25 (P60880), Rab-interacting lysosomal protein (Q96NA2), α-synuclein (P37840), 4E-BP1 (Q13541), 4E-BP2 (Q13542), NCoA3 (Q9Y6Q9), cyclin-dependent kinase inhibitor 1B or p27 (P46527), Cbp/p300-interacting transactivator 2 or CITED2 (Q99967), hypoxia-inducible factor 1-alpha or HIF-1alpha (Q16665), and the cystic fibrosis transmembrane conductance regulator (P13569). The Protein Data Bank (PDB, https://www.rcsb.org/) IDs that were used in this work are: 4E-BP2 (3am7, 2mx4), 4E-BP1 (5bxv), NCoA3/ACTR (1kbh), p27 (1jsu), SNAP25 (1kil, 1xtg), HIF-1alpha (1l8c), CITED2 (1p4q), and CFTR (1r0x, 1xmi). BMRB accession codes 6968 (α-synuclein), 5744 (α-synuclein bound to SDS micelles), 19114 (4E-BP2), 19905 (phosphorylated 4E-BP2), 15397 (ACTR), and 5228 (ACTR bound to CBP) were downloaded from the BMRB (https://bmrb.io/).

## Code Availability

The code and data used in this paper are available on GitHub (https://github.com/IPritisanac/AF2.IDR).

## Competing Interests

The authors do not have any competing interests.

## Acknowledgements

We thank Dr. Adriaan Bax (National Institutes of Health, USA) and William Ford Freyberg (University of Wisconsin-Madison, USA) for critical comments and feedback on the manuscript, Dr. Giuliana Fusco (University of Cambridge, UK) for sharing the experimental CD spectrum of α-synuclein, Dr. Kresten Lindorff-Larsen (University of Copenhagen, Denmark) for making the α-synuclein SAXS data available via GitHub, and Emil Spreitzer and Dr. Tobias Madl (Medical University of Graz, Austria) for sharing the solvent PRE data from α-synuclein. This study made use of NMRbox: National Center for Biomolecular NMR Data Processing and Analysis, a Biomedical Technology Research Resource (BTRR), which is supported by NIH grant P41GM111135 (NIGMS). TRA and IP were supported by a Banting Postdoctoral Fellowship from the Canadian Institutes of Health Research (CIHR) and a LiUNA! Fellowship for Research Innovation from The Hospital for Sick Children, respectively. AMM and JDF-K acknowledge support from the CIHR (CIHR Foundation Grant (grant no. FDN-148375) to JDF-K; CIHR grant no. PJT-148532 to AMM and JDF-K) and the Canada Foundation for Innovation (CFI) for funding to AMM. JDF-K holds a Canada Research Chair in Intrinsically Disordered Proteins.

## Notes

### Competing Interest Statement

The authors have declared no competing interest.

### Summary of Updates

New analyses and data presented in Figure 5, Figure 7, and Figure 8. The Supplemental Information has also been updated with new analyses, and a new author was involved (Desika Kolaric).

