## Supporting Information for "Systematic identification of conditionally folded intrinsically disordered regions by AlphaFold2"

[§] Equal contribution

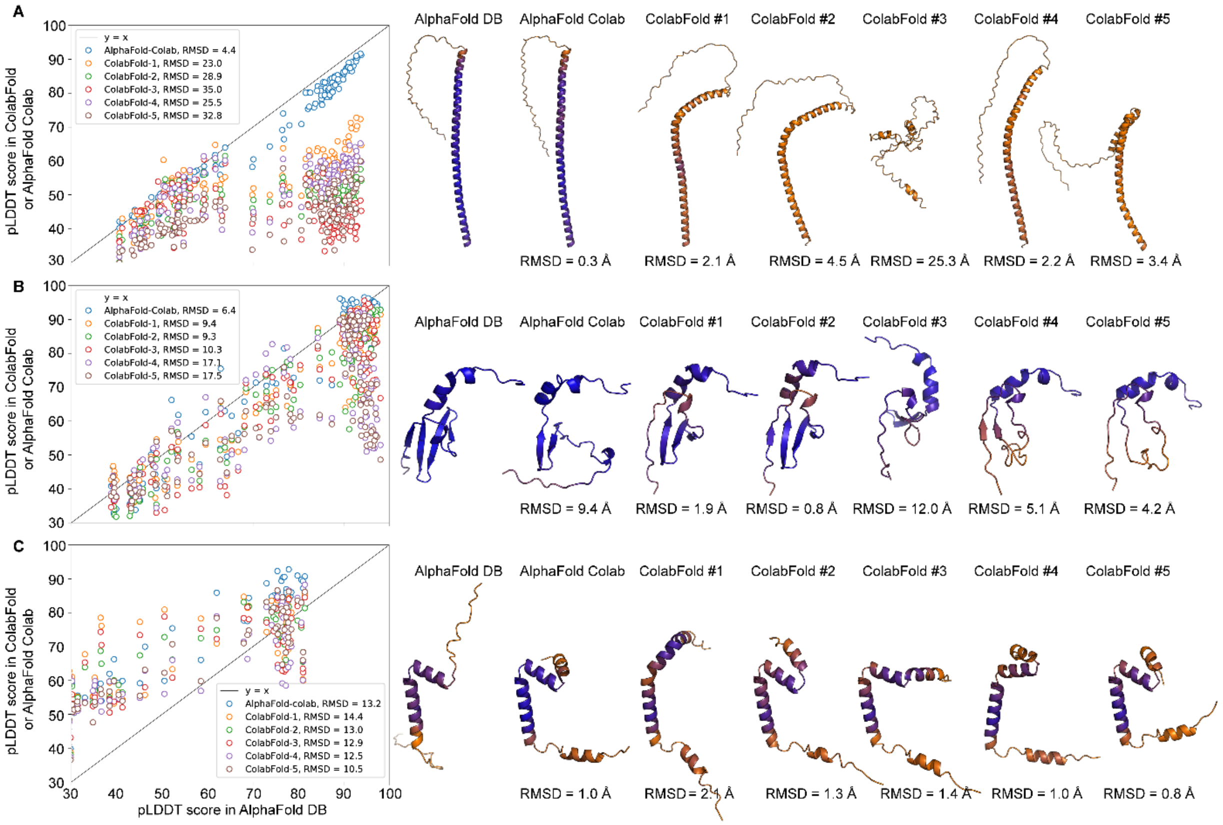


**Supplementary Figure 1. Comparison of the structural predictions by the AFDB and other online versions of AlphaFold2.** ColabFold (Mirdita et al. 2021) and the AlphaFold Colab notebooks were used to generate structural predictions of human α-synuclein (UniProt: P37840), 4E-BP2 (Q13542), and ACTR (Q9Y6Q9; residues 1123-1193). The per-residue pLDDT scores were extracted from the resultant structure (AlphaFold Colab) or the five models produced by ColabFold and compared to those within the AFDB structure. The root-mean-squared-deviation (RMSD) of the pLDDT scores is listed for the comparison of AFDB with AlphaFold Colab or ColabFold. *Right*: the structures generated from the AFDB, AlphaFold Colab, and ColabFold calculations. Note that AlphaFold Colab returns only one model whereas ColabFold returns five models. The RMSD values over regions of secondary structure upon alignment to the AFDB model are listed.

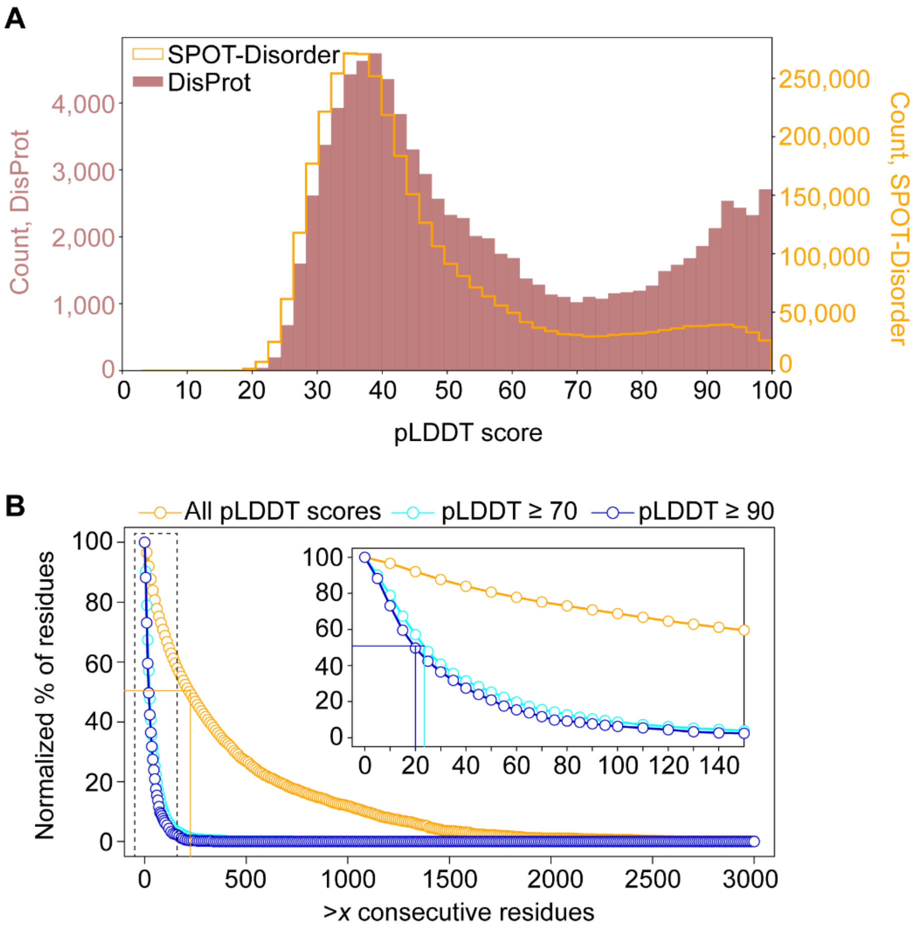


**Supplementary Figure 2**. **Filtering the human AFDB with DisProt and for regions of consecutive disorder.** (**A**) Histogram of per-residue pLDDT scores for proteins in the human AFDB filtered by SPOT-Disorder-predicted regions of intrinsic disorder (orange) or experimentally validated IDRs from DisProt (maroon). The percentage of residues with pLDDT scores ≥ 70 is 14.3% for SPOT-Disorder and 29.5% for DisProt. (**B**) SPOT-Disorder-predicted disordered residues (orange) were filtered for regions that contained greater than *x* consecutive residues that were disordered. Long disordered regions are abundant in the human proteome, with over 50% of SPOT-Disorder-predicted disordered residues falling in regions that have more than 220 consecutive residues (orange line). When only SPOT-Disorder-predicted IDRs that have pLDDT scores ≥ 70 (cyan) or ≥ 90 (blue) are analyzed, the percentage of residues that have long, consecutive regions of disorder is dramatically smaller, with 50% of residues in these groups of IDRs having more than 24 (cyan line) or 20 (blue line) consecutive residues for pLDDT scores ≥ 70 and ≥ 90, respectively. The y-axis shows the normalized percentage of residues, *i.e.* the sum of the consecutive residues for each threshold divided by the total number of residues in each group of IDRs.

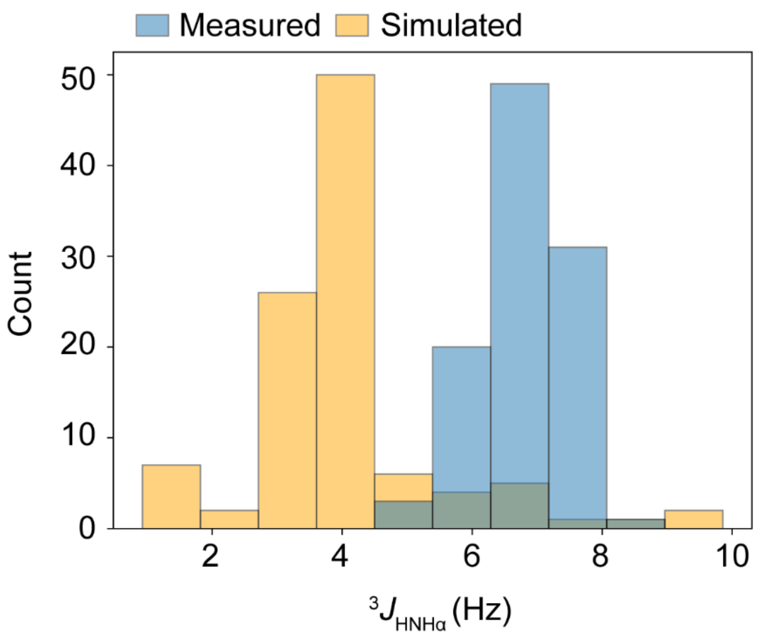


**Supplementary Figure 3**. **Using NMR data without chemical shift assignments to assess an AlphaFold2 prediction**. $^3J_{HNH\alpha}$ coupling constants (Mantsyzov et al. 2014) from α-synuclein (blue) are shown in a histogram format. The histogram of $^3J_{HNH\alpha}$ values that are back-calculated from the AlphaFold2 structural model are shown in orange. This histogram approach for NMR values mimics a situation in which resonance assignments are not available. A comparison between the histograms shows clear discrepancies between the experiment and simulation, even when no assignments are available.

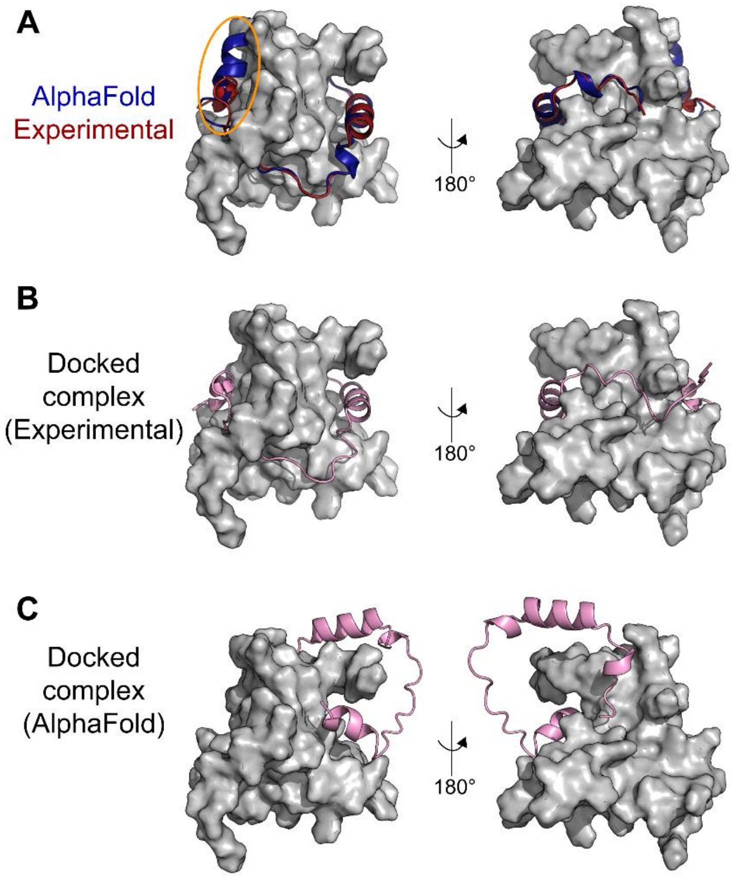


**Supplementary Figure 4. The structures of IDRs in the AFDB, even if confident, may yield incorrect results when used to obtain a structural model of a protein complex using molecular docking**. (**A**) Overlaid structures of the CITED2 TAD bound to CBP TAZ1 that was experimentally determined (red, PDB 1p4q) and the AlphaFold-predicted structure of the CITED2 TAD (blue). The TAZ1 domain is shown in a grey surface representation and was not included in the AlphaFold prediction of the structure for the CITED2 TAD. The orange ellipse indicates the position of the CITED2 C-terminal helix whose orientation differs in the AlphaFold model with respect to the experimental structure. The lowest-energy model returned by FRODOCK2.0 (Ramírez-Aportela et al. 2016) when docking the CITED2 TAD with the (**B**) experimentally determined structure or the (**C**) AlphaFold-predicted structure onto the CBP TAZ1. Note that the C-terminal helix of CITED2 does not interact with the correct binding pocket in the CBP TAZ1 domain when using the AlphaFold model for docking purposes. See text for residue boundaries.

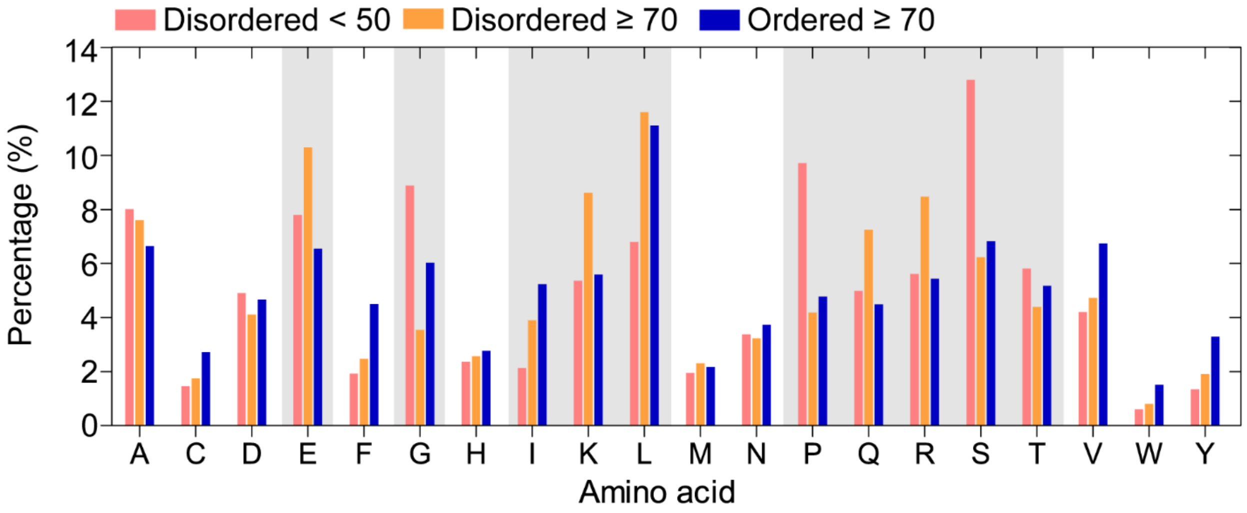


**Supplementary Figure 5. Amino-acid frequencies in the human AFDB as a function of pLDDT score.** The human AFDB was filtered with SPOT-Disorder-predicted regions of disorder to obtain predicted disordered and ordered regions. Plotted here are amino-acid percentages (eq 1, Methods) in predicted disordered regions with pLDDT scores < 50 (salmon), predicted disordered regions with pLDDT scores ≥ 70 (orange), or predicted ordered regions (blue). Amino acids with particularly significant differences between the two disordered datasets are indicated with grey boxes. These raw percentages were used to compute the amino acid differences shown in Figure 5.

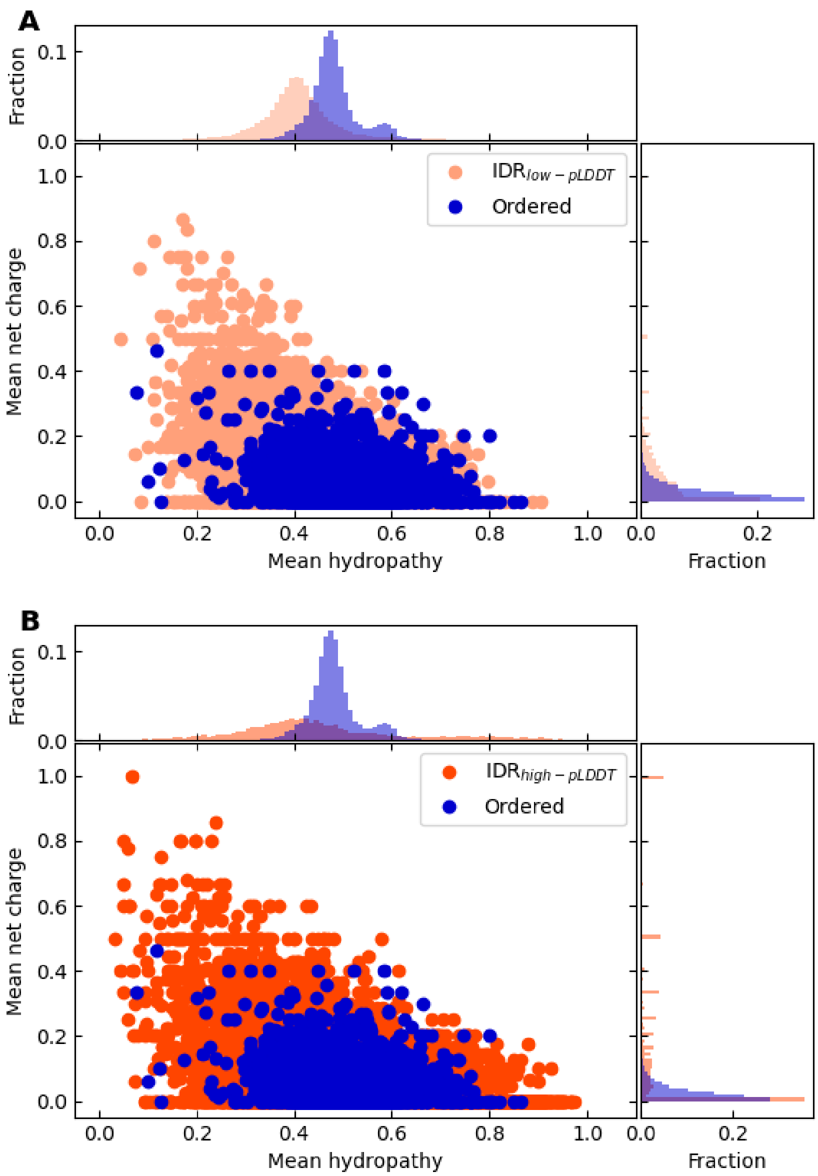


**Supplementary Figure 6. Predicted IDRs in the human AFDB with high-confidence structures resemble IDR sequences.** Analysis of the mean net charge and mean hydropathy values of predicted IDRs that have very low (≤ 50, $IDR_{low\text{-}pLDDT}$) (**A**) or very confident (**B**) (≥ 90, $IDR_{high\text{-}pLDDT}$) pLDDT scores as compared to ordered regions. The histograms are normalized such that the sum of all values equals unity.

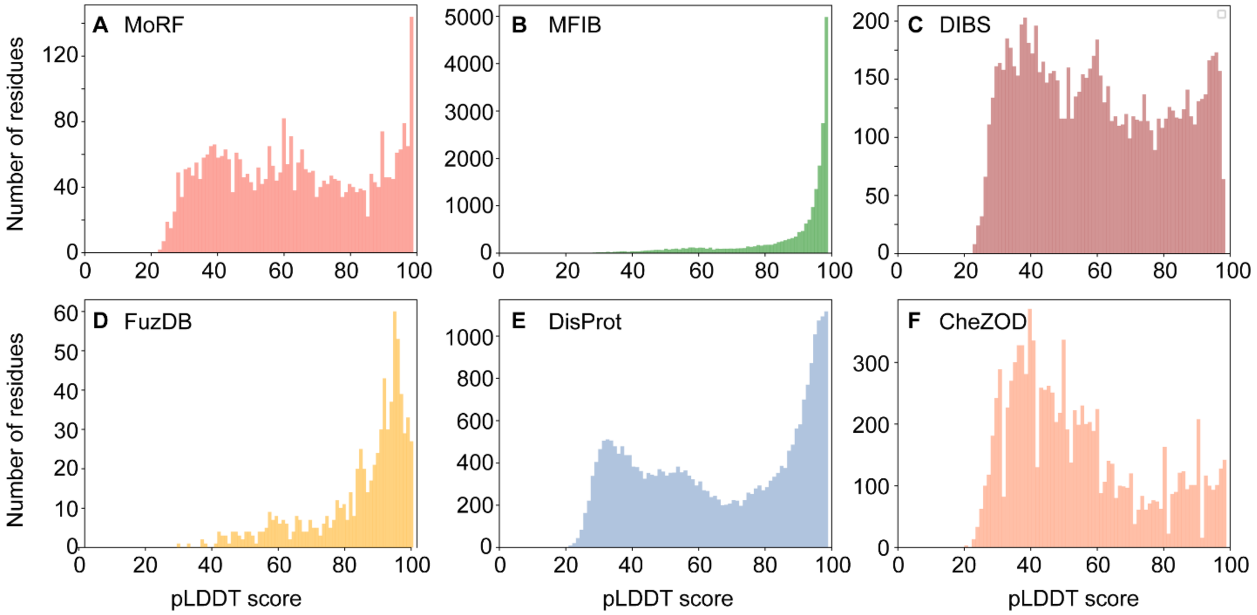


**Supplementary Figure 7. Per-residue pLDDT scores for IDRs that were used to classify conditional folders**. Analysis of per-residue pLDDT scores for known IDRs/IDPs that fold upon binding or modification. These examples were taken from the following five databases: MoRF (**A**), MFIB (**B**), DIBS (**C**), FuzDB (**D**), and DisProt (**E**). Of the IDR sequences in these databases, only the regions that map to the corresponding predicted structure in the AFDB were retained for this analysis. See main text for details. The y-axis shows the counts for the pLDDT scores of the IDRs/IDPs in the indicated databases. Panels **A** – **E** were used as true positive datasets in the ROC analysis. For comparison, the pLDDT score distribution from the true negative dataset (CheZOD, filtered as described in the main text), is shown in (**F**). The IDRs in the filtered CheZOD dataset have not been reported to conditionally fold. Note that the pLDDT score distribution for these IDRs is shifted toward lower values.

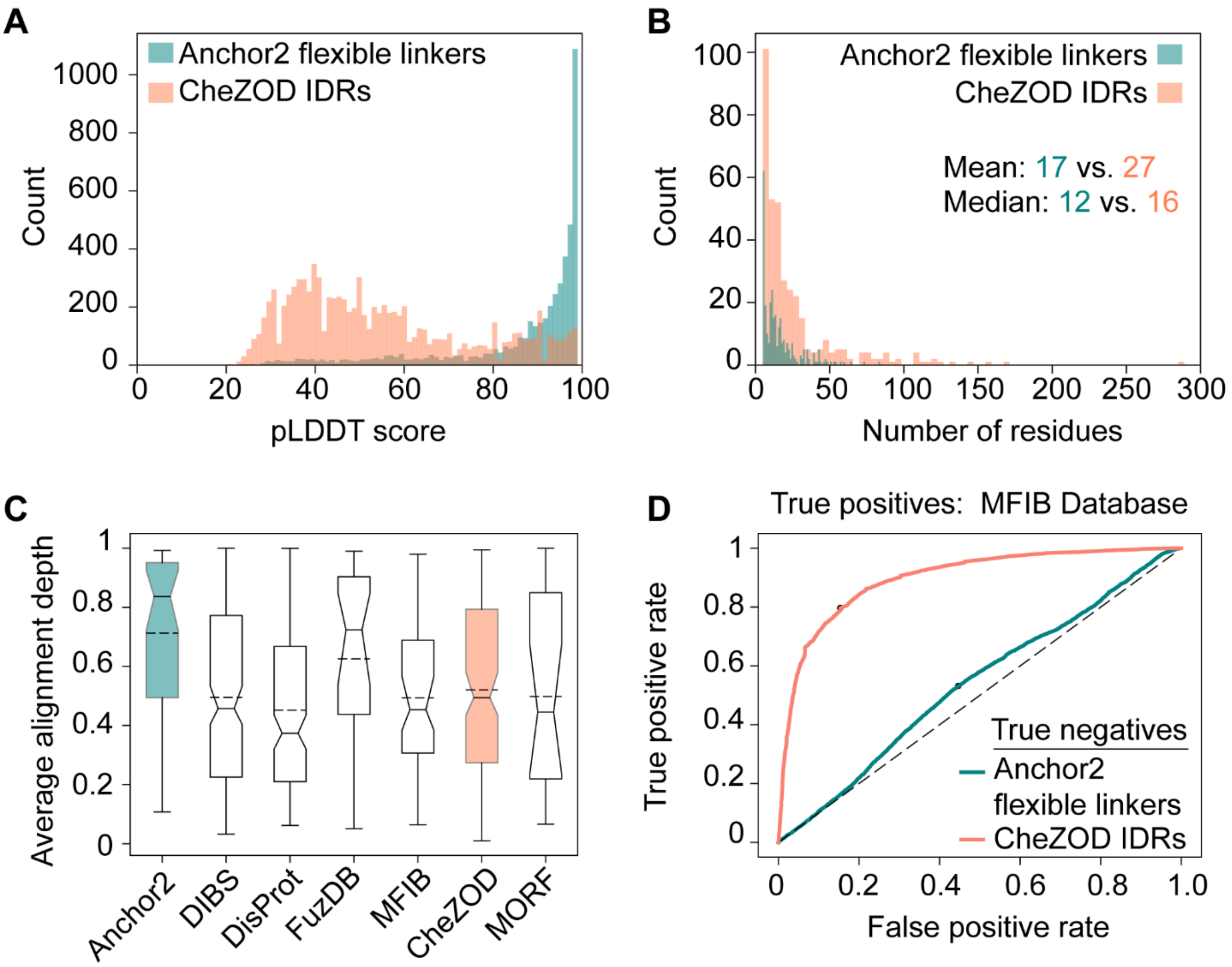


**Supplementary Figure 8**. **Comparison of NMR-validated IDRs from CheZOD with the dataset of flexible linkers used by Anchor2.** (**A**) Distribution of per-residue pLDDT scores for the IDR sequences in the Anchor2 true negative dataset of flexible linkers (blue). Per-residue pLDDT scores for the CheZOD-filtered set of IDRs that do not conditionally fold are shown in orange. The distribution of pLDDT scores for the filtered CheZOD dataset more closely resembles IDRs in the human proteome (Figure 1). (**B**) The distribution of IDR sequences in the Anchor2 flexible linkers dataset (blue) and the CheZOD-filtered IDR dataset (orange). The mean and median values for each dataset are indicated (Anchor2 vs. CheZOD). (**C**) Box plots showing the average alignment depth for the IDR sequences in the Anchor2 flexible linker dataset (blue) as compared to the filtered CheZOD dataset (orange). For comparison, the average alignment depths are shown for the five true positive databases of IDRs that are known to conditionally fold (DIBS, DisProt, FuzDB, MFIB, MORF). The IDRs in the flexible linkers dataset from Anchor2 have the deepest sequence alignments. (**D**) ROC curves for the binary classification task of identifying conditionally folded IDRs based on pLDDT scores alone. The true positive dataset was the MFIB database and the true negatives were either the Anchor2 flexible linkers (blue) or the filtered set of CheZOD IDRs (orange). The classification performance greatly increases when the CheZOD-derived IDRs are used as a set of true negatives.

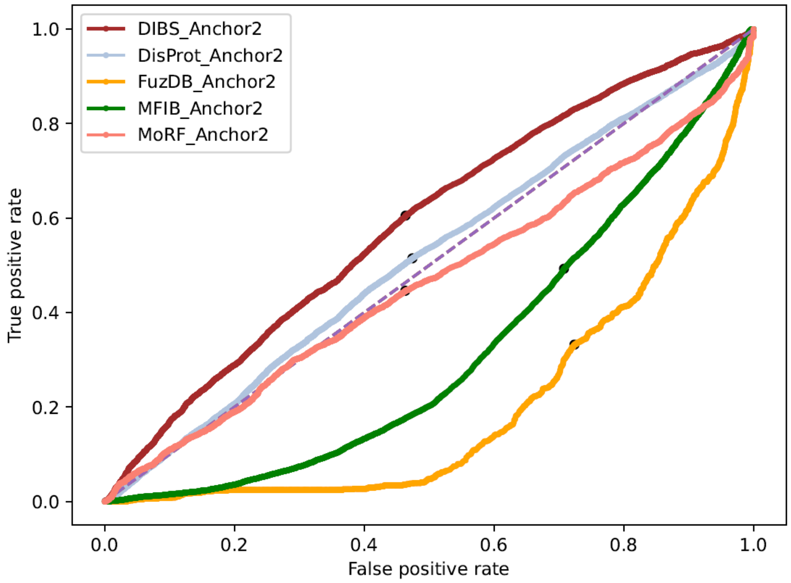


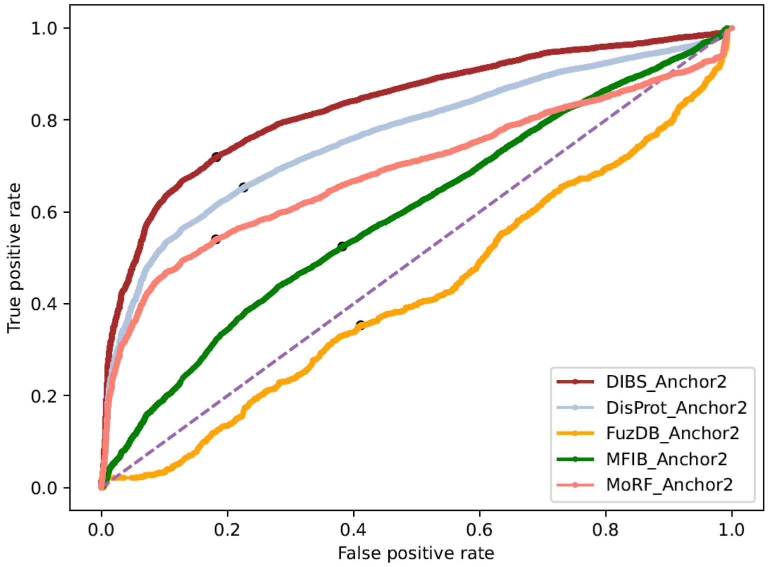


**Supplementary Figure 9. Performance of the Anchor2 software as a classifier of conditionally folding IDRs.** (*Top*) Anchor2 was run with default parameters using the sequences of IDRs from five databases of conditionally folded IDRs (MFIB, DIBS, DisProt, FuzDB, MoRF) as input. The set of true negatives was the filtered set of NMR-validated IDRs in CheZOD (see Supplementary Figure 8). An ROC analysis was performed on these data. Anchor2 performs well at classifying the IDRs within the DIBS dataset, although some of these IDRs were used in the training of the Anchor2 software. The ROC curves when using AlphaFold2 pLDDT scores as input data are shown in Figure 6A for comparison. (*Bottom*) The same analysis but with the flexible linkers dataset supplied as true negatives (see text, Supplementary Figure 8), which was used for the training and testing of Anchor2. The AUC for DIBS is 0.83 and closely recapitulates the original Anchor2 performance (Mészáros et al. 2018).

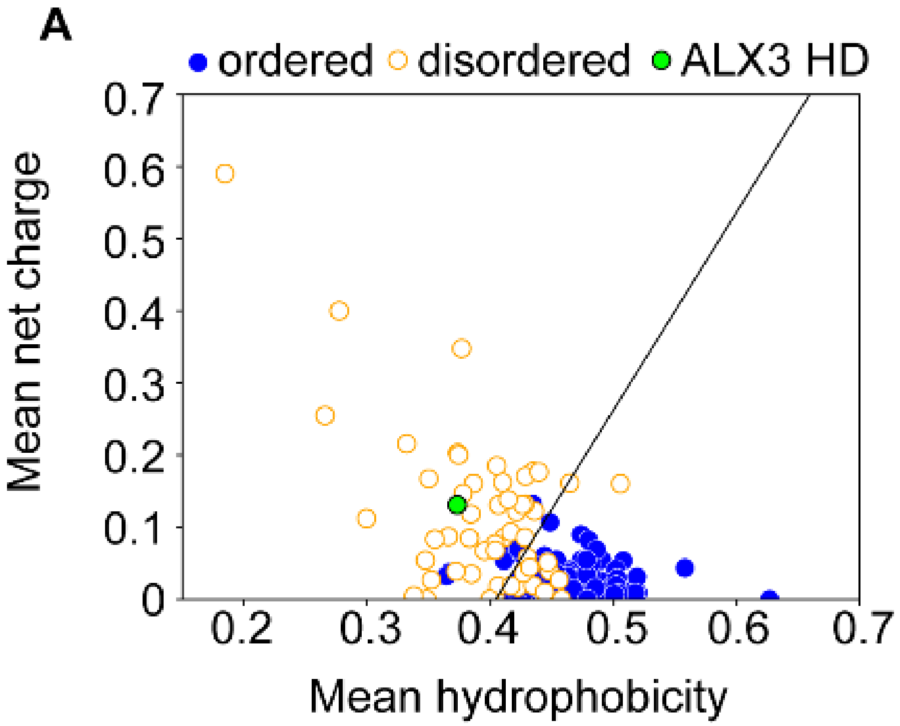


**Supplementary Figure 10. Sequence analysis of ALX3.** (**A**) Uverksy plot showing ordered (blue) and disordered (orange) proteins separated by the mean hydrophobicity and mean net charge of the sequence. The ALX3 homeodomain (green), residues 153-213, resembles the sequence properties of disordered proteins. The ordered and disordered proteins were extracted from the PONDR website (Romero et al. 2001).

**Supplementary Table 1**. **Segmenting the AFDB into predicted regions of disorder and order**. SPOT-Disorder and IUPred2A were used to predict IDRs within the human proteome and then to segment the AFDB into ordered and disordered regions. The number of residues that are not mapped from the human proteome to the AFDB are listed in the final column. The percentage of the predicted ordered, predicted disordered, and not mapped residues are listed in parentheses.

| Dataset used to filter AFDB | Total residues | Ordered (%) | Disordered (%) | Not mapped (%) |
|---|---|---|---|---|
| SPOT-Disorder | 10,825,508 | 7,127,685 (65.84%) | 3,539,799 (32.70%) | 158,024 (1.46%) |
| IUPred2A | 10,825,508 | 7,942,769 (73.37%) | 2,714,596 (25.08%) | 168,143 (1.53%) |

**Supplementary Table 2. Per-residue pLDDT scores for predicted disordered and ordered regions of the human proteome.** SPOT-Disorder was used to predict disordered regions of the human proteome and then to segment the human AFDB into regions of predicted disorder and order. The percentage of residues are listed as a function of the per-residue pLDDT score. Values listed in parentheses were obtained when IUPred2A was used to predict regions of disorder.

| Dataset | *N* residues | % pLDDT ≥ 90 | % 90 > pLDDT ≥ 70 | % 70 > pLDDT ≥ 50 | % pLDDT < 50 |
|---|---|---|---|---|---|
| Proteome | 10,825,508 | 38.12 | 24.46 | 9.85 | 27.57 |
| Ordered | 7,127,685 | 54.51 | 31.82 | 7.46 | 6.21 |
| Disordered | 3,539,799 | 4.52 (7.39) | 9.78% (10.36) | 14.73 (12.88) | 70.98 (69.37) |

**Supplementary Table 3. SPOT-Disorder-predicted IDRs in the human proteome that have high- or very high-confidence AlphaFold2 pLDDT scores.** SPOT-Disordered-predicted IDRs in the human proteome of a minimum length of 10 or 30 or more consecutive residues are quantified. The pLDDT score is also considered, with any pLDDT score (*all*) or only residues with pLDDT scores greater than or equal to 70 or 90. *N* IDRs refers to the number of regions, *N* proteins refers to the number of unique proteins (*i.e.*, multiple IDRs may come from a single protein), and *N* residues to the total number of amino acids. The superscripts [a] and [b] indicate the number of residues in each pLDDT threshold divided by the total number of disordered residues for a length cut-off of ≥ 10 and ≥ 30, respectively.

| pLDDT score | Length cut-off | *N* proteins | *N* IDRs | *N* residues |
|---|---|---|---|---|
| all | ≥ 10 AA | 17,333 | 37,616 | 3,398,344 |
| ≥ 70 | ≥ 10 AA | 7,586 | 14,996 | 400,244 (11.8%)[a] |
| ≥ 90 | ≥ 10 AA | 2,981 | 4,883 | 121,314 (3.6%)[a] |
| all | ≥ 30 AA | 13,070 | 20,492 | 3,097,338 |
| ≥ 70 | ≥ 30 AA | 2,409 | 3,730 | 210,584 (6.8%)[b] |
| ≥ 90 | ≥ 30 AA | 843 | 1,157 | 61,745 (2%)[b] |

**Supplementary Table 4. Secondary structure content in the predicted disordered and ordered regions as a function of the pLDDT score.** SPOT-Disorder was used to segment the human AFDB into predicted regions of disorder and order.

| Dataset | pLDDT score | Helix (%) | Coil (%) | Strand (%) |
|---|---|---|---|---|
| Ordered | ≥ 90 | 47.2% | 28.5% | 24.3% |
| | 90 > *x* ≥ 70 | 44.0% | 42.6% | 13.4% |
| | 70 > *x* ≥ 50 | 33.2% | 61.7% | 5.1% |
| | < 50 | 10.5% | 87.8% | 1.7% |
| Disordered | ≥ 90 | 78.1% | 16.1% | 5.8% |
| | 90 > *x* ≥ 70 | 66.5% | 29.0% | 4.5% |
| | 70 > *x* ≥ 50 | 30.8% | 67.4% | 1.8% |
| | < 50 | 3.4% | 96.4% | 0.2% |

**Supplementary Table 5. Classification performance statistics from the ROC analysis of conditionally folded IDRs.** Listed here are the AUC, recall, precision, and PPV values from the ROC analysis of AlphaFold2 per-residue pLDDT scores in classifying conditionally folded IDRs. The true positive databases are listed under "Database name" and the set of true negatives was the filtered CheZOD list.

| Database name | AUC | RECALL | PRECISION | PPV |
|---|---|---|---|---|
| MFIB | 0.90 | 0.80 | 0.92 | 0.84 |
| FuzDB | 0.85 | 0.81 | 0.23 | 0.79 |
| DisProt | 0.64 | 0.55 | 0.84 | 0.65 |
| MoRF | 0.61 | 0.55 | 0.37 | 0.62 |
| DIBS | 0.58 | 0.52 | 0.58 | 0.58 |

### Supplementary Appendix

In order to rapidly determine if the AlphaFold structure of an IDR/IDP is accurate, it is necessary to collect biophysical data on a purified sample of the IDR/IDP of interest. While biophysical experiments that can be performed inside living cells or in cell lysate may also be applicable (Gronenborn & Clore 1996; Gruebele & Pielak 2021), and do not require sample purification, we focus here on *in vitro* measurements performed on purified proteins as such experiments are performed more routinely. Moreover, it would be beneficial to minimize the time and cost associated with sample preparation, experimental acquisition, and data analysis. Therefore, we have concentrated on biophysical assays that can be performed on (1) the wild-type amino-acid sequence of the protein, *i.e.* there should be no mutations required for covalent linkage of fluorescent tags or spin labels, and (2) natural abundance protein in a standard buffer, *i.e.* there should be no need for isotope enrichment or $D_2O$-based buffers that are common in small-angle neutron scattering (SANS), NMR, and EPR (Clifton et al. 2019; Gardner & Kay 1998; Schmidt et al. 2016). As an example, we have compared biophysical data recorded on the protein α-synuclein, for which a number of biophysical measurements are available, with those that were simulated from the AFDB structure (**Figure 5**, **Supplementary Figure 3**).

A global measurement of secondary structure content from circular dichroism (CD) or NMR spectroscopy may prove valuable to quickly distinguish between plausible models. In the latter case, neither resonance assignments nor isotope-enrichment with $^{15}N$ or $^{13}C$ are needed for global secondary structure analysis by NMR (Wishart et al. 1991). However, the NMR approach requires relatively high protein concentrations, or long acquisition times in case of limited sample amounts, and a skilled user to analyze the data. For CD spectroscopy, sample requirements are minimal, data can be acquired in a short time, and one can quickly identify the secondary structure with software programs that fit the experimental spectrum (Greenfield 2006). The experimental CD spectrum can readily be compared to simulated CD spectra obtained from an input three-dimensional structure via the webserver PDB2CD (Mavridis & Janes 2017). For example, the predicted CD spectrum of α-synuclein based on the structure in the AFDB has diagnostic minima at 208 and 222 nm, characteristic of helical secondary structure, whereas the experimental spectrum shows a minimum near 200 nm, which is indicative of random coil conformations (**Figure 5A**). These features show that the experimental data reflect a largely random or statistical coil conformation whereas the spectrum of the AFDB structure would be dominated by signals from α-helical conformations.

In cases where global secondary structure quantification is insufficient to determine if the AFDB model is accurate, such as when the AlphaFold structure of an IDR may only have a small percentage of secondary structure, measurements that are sensitive to the shape of the molecule may provide more information content. For example, it has already been demonstrated that IDRs with low pLDDT scores can be too expanded relative to experimental measurements (Ruff & Pappu 2021). Thus, dynamic light scattering (DLS), small-angle X-ray scattering (SAXS), and pulsed-field gradient diffusion NMR spectroscopy (PFG-NMR), which are all highly sensitive to the shape of a molecule, could quickly identify such structures as erroneous. Moreover, all of these experiments can be performed on label-free, natural-abundance protein samples in conventional $H_2O$-based buffers. The hydrodynamic properties that are sampled in DLS and PFG-NMR experiments can be simulated from a known structure using the HYDROPRO software package (Ortega et al. 2011), while SAXS curves can be simulated from a known structure with Crysol (Svergun et al. 1995). We have chosen here to focus on PFG-NMR and SAXS, because DLS is more sensitive for larger particles and may require high protein concentrations for shorter IDRs in order to achieve sufficiently high signal-to-noise. By contrast, both PFG-NMR and SAXS data can be collected on relatively short IDRs at low protein concentrations. Moreover, because IDRs typically tumble independently of globular domains and yield sharp signals, NMR can monitor IDRs within very large particles even when the signals from the globular domain cannot be detected (Carver et al. 1992).

The simulated PFG-NMR and SAXS data for α-synuclein (**Figure 5B, 5C**). The PFG-NMR data show that α-synuclein diffuses faster than the expectation based on the AFDB structure (**Figure 5B**), and the SAXS data show deviations in the low *q* regions (**Figure 5C**). The fitted values of the radius of gyration ($R_g$) and maximum distance ($D_{max}$) are 35.6 ± 0.2 Å and 109 Å, whereas the back-calculated values from the AFDB structure are 42.6 Å and 152 Å, respectively. Thus, both PFG-NMR and SAXS measurements would indicate that the AFDB model is too extended relative to the conformation in solution.

If global measurements are insufficient to determine if the AFDB model is accurate, then more detailed local structural parameters can be obtained from NMR spectroscopy (Wishart et al. 1991). Such experiments, however, generally require resonance assignment in order to yield atomic-level information. The process of resonance assignment typically requires the preparation of $^{13}C$,$^{15}N$-labeled protein and the collection of multiple sets of 3D NMR spectra, which requires considerable more time than the above-mentioned experiments. Therefore, the following experiments can no longer be classified as “rapid”; however, the information content afforded by these experiments is very high.

For example, the deviation of the measured $^{13}C\alpha$ chemical shifts from random coil values (secondary chemical shift) is highly sensitive to α-helical conformations (**Figure 5D**). For β-strands, $^{13}C\beta$ shifts are more sensitive. A comparison of secondary $^{13}C\alpha$ chemical shifts for α-synuclein and those back-calculated from the AFDB structure with SPARTA+ (Shen & Bax 2010) reveals strong α-helical conformations in the AFDB model that are not present in the measured data. The residue-specificity afforded by NMR further pinpoints the structural deviations to the first *ca.* 90 residues of the AFDB model. In addition, three-bond scalar couplings report on the intervening dihedral angles. In particular, the readily measurable $^{3}J_{HNH\alpha}$ coupling reports on the dihedral angle Φ. For α-synuclein, the measured $^{3}J_{HNH\alpha}$ coupling constants (Mantsyzov et al. 2014) are considerably more uniform than those back-calculated from the AFDB model using the parametrized Karplus equation (Methods, equation 3) (Vögeli et al. 2007) (**Figure 5E**). As above, the residue-specificity of these measurements enables a site-by-site comparison of the coupling constants. Finally, solvent paramagnetic relaxation enhancements (sPREs) provide site-specific structural restraints that report on solvent accessibility and local structural conformation. The measured $^{1}H\alpha$ sPREs (Hartlmüller et al. 2019) are considerably larger for the first *ca.* 90 residues of α-synuclein than those back-calculated from the AFDB structure with sPRE-calc (Gong et al. 2017) (**Figure 5F**). Collectively, the NMR experiments point to localized structural differences in the first *ca.* 90 residues of the AFDB structure relative to the conformation sampled in solution. Additional NMR experiments, such as residual dipolar couplings (RDCs), PREs with site-directed spin labelling, and $^{15}N$ relaxation provide further orientational and distance restraints that can be compared to a structural model.

Thus, relatively easy biophysical experiments (**Figure 5A-C**) can be performed on purified proteins *in vitro* to quickly assess the accuracy of AFDB predictions. These experiments do not require the introduction of any labels or mutations and can be performed in standard buffers that are compatible with conventional biophysical and biochemical assays. More time-consuming NMR experiments (**Figure 5D-F**) can be performed to obtain site-specific information.
